# Lamins gate nuclear and chromatin structures for cardiomyocyte maturation genes

**DOI:** 10.64898/2026.05.20.726565

**Authors:** Katherine A. Bossone, Xiaobin Zheng, Lidya Kristiani, Reni Marsela, Youngjo Kim, Yixian Zheng

## Abstract

The nuclear lamina contains intermediate filaments, called lamins, which function to maintain nuclear integrity and organize the <u>L</u>amina-<u>A</u>ssociated chromatin <u>D</u>omains (LADs). Despite near ubiquitous expression, lamins show cell-type-specific functions. In the heart, they support epicardial cell migration, cardiomyocyte nuclear integrity and maturation. How these functions are integrated with different gene expression programs in these cells during heart development is unknown. We show that cardiomyocytes require lamin-A and -B1 for perinatal mouse survival. Importantly, lamin-B1 facilitates timely cardiomyocyte maturation by maintaining LADs and chromosome territories. By examining changes in cardiomyocyte gene expression and 3D genome organization upon lamin-B1 deletion, we show lamin-B1 maintains chromatin neighborhoods, which can in turn support transcription factors that regulate genes involved in cardiomyocyte structural maturation and gradual cessation of cell division. We propose a genomic logic by which the widely expressed lamins collaborate with transcription factors to specifically promote cardiomyocyte maturation and cell cycle exit during development.

**Highlights:**

1. Lamin-A/B1 dose-dependently set cardiomyocyte nuclear order and postnatal survival.
2. Lamin-B1 loss disrupts transcription programs for cardiomyocyte maturation.
3. Lamin-B1 shapes chromatin neighborhoods to support cardiomyocyte maturation programs.

## Introduction

Organism development involves lineage specification and cell proliferation, followed by terminal differentiation and cell cycle exit in different cell lineages^1,2^. These processes are regulated by different transcriptional programs. The timely execution of these transcription programs depends not only on lineage-specific transcription factors but also on three-dimensional (3D) genome organization that partitions regulatory elements into spatially and functionally distinct nuclear environments^3^.

Based on 3D chromatin interaction maps, the genome can be partitioned into the gene-rich, transcriptionally active A compartment and the gene-poor, transcriptionally repressive B compartment^3–5^. Within these compartments are megabase to sub-megabase sized <u>T</u>opologically <u>A</u>ssociating chromatin <u>D</u>omains (TADs) that are established by CTCF and cohesin proteins. A role of TADs is to constrain enhancer-promoter contacts and help shape tissue-specific transcription programs^6,7^. The B compartment is also largely composed of Lamina Associated Domains (LADs), which are heterochromatic genome regions with overall low gene expression^8,9^. Previous work showed evidence of LADs remodeling during lineage development^10–12^, and some nuclear lamina proteins are known to support organismal development^13,14^. It remains unclear how the nuclear lamina coordinates transcriptional programs to support development, especially in cell maturation and cell division.

The nuclear lamina contains ∼100 proteins including lamins, inner nuclear envelope proteins, and chromatin binding proteins^15^. The nuclear lamina interacts with the nuclear pore complexes (NPCs)^16–20^. Lamins ensure even NPC distribution around the nuclear envelope^16–19^, while NPC basket components coordinate mRNA export^21,22^, and recruitment of transcription factors and chromatin regulators^23^. The nuclear lamina also interacts with another nuclear envelope structure called the Linker of Nucleoskeleton and Cytoskeleton (LINC) complex, consisting of the SUN- and KASH-domain proteins in the inner and outer nuclear envelopes, respectively^24^. Lamins interact with the Sun1 protein, and ensure an even distribution of the LINC complex^18^. Studies suggest that these interactions anchors the nuclear lamina to the cytoskeleton and transmits mechanical cues that feed into signaling cascades controlling transcription factors and cell-cycle regulators^25,26^. It is believed that these complicated physical and functional connections between the nuclear lamina, NPCs, and LINC complexes can gate signaling through NPCs, and couple mechanotransduction to transcriptional regulation and cell-cycle decisions. How these connections direct a given cell type to make cell-cycle decisions during terminal differentiation remains unclear.

Studies show the loss or mutation of nuclear lamina proteins, such as lamins, frequently produce nuclear envelope ruptures^27–29^, chromatin protrusions^27^, and replication-associated DNA damage^30^. Secondary effects on genome integrity and p53 activation can confound interpretation of the roles in transcription and differentiation^31^. Nevertheless, extensive research has shown lamins are required for proper animal development^13,14^. Lamin-A mutations cause human “laminopathies” including muscular dystrophy, lipodystrophy, Hutchinson-Gilford progeria, and dilated cardiomyopathy^32^. Perturbation of different lamins in different model organisms and stem cells disrupt myogenesis, adipogenesis, neurogenesis, and germline differentiation^14,33–35,17^. Despite the clear developmental functions, how these lamins collaborate with developmental transcriptional programs has been difficult to establish.

Cardiomyocyte development offers an attractive model for probing how lamins couple transcriptional maturation and cell-cycle exit because cardiomyocytes undergo a clearly defined stereotypical transition into maturation. This is characterized by mononucleated, proliferative cardiomyocytes becoming binucleated, polyploid cells that permanently exit the cell cycle while simultaneously activating structural and metabolic maturation programs^36^. This tight temporal developmental coupling offers a great opportunity to examine whether and how lamins collaborate with transcription factors to simultaneously drive maturation and halt cell division.

Mammals express three lamin genes, *Lmna* (encoding lamin-A or lamin-C through alternative splicing), *Lmnb1* (encoding lamin-B1), and *Lmnb2* (encoding lamin-B2). The B-type lamins are expressed throughout embryogenesis in most tissues, whereas lamin-A/C expression gradually increases during embryonic and/or postnatal development in many cell lineages including kidney cells, certain neuronal subtypes of the retina, skeletal muscle, and cardiomyocytes^14,15,37–39^. In the heart, *Lmna* mutations are among the most common monogenic causes of dilated cardiomyopathy (DCM) with conduction disease^40^. Cardiomyocyte-specific deletion of *Lmna* in mice causes severe, early-onset DCM, nuclear rupture, severe DNA damage, and lethality within 3-4 weeks after birth^41^. While lamin-B2 is expressed throughout heart development, its expression in cardiomyocytes ends shortly after birth and coincides with cardiomyocyte cell cycle exit. Cardiomyocyte-specific deletion of lamin-B2 during early embryonic development leads to impaired cardiac regeneration, increased nuclear polyploidy, and defects in cell division^42^. Therefore, lamin-B2 appears to support cardiomyocyte proliferation during development. The cardiomyocyte-specific role of lamin-B1 remains unknown, despite its constitutive expression^15^.

By analyzing the cardiomyocyte-specific roles of lamin-A and -B1 in the settings of single knockout, single knockout with haplo-insufficiency, and double knockout conditions, we discover a synergistic role for lamin-B1 and lamin-A in supporting cardiomyocyte development and mouse survival. Since cardiomyocyte-specific loss of lamin-B1 does not cause DNA damage, we are able to show that this lamin regulates cardiomyocyte structural and metabolic maturation, and cell cycle exit. By analyzing how lamin-B1 supports cardiomyocyte chromosome territories, LADs, 3D genome organization, and gene expression, we show that lamin-B1 can collaborate with transcription factors known to regulate cardiomyocyte maturation programs to support their maturation and cell cycle exit.

## Results

### Lamin-A and lamin-B1 loss in cardiomyocytes causes severe heart defects and early postnatal mouse lethality

Studies in mice have shown that lamin-A expression is low in cardiomyocytes during embryogenesis^39^ and its specific deletion in cardiomyocytes during early embryonic development using αMyHC-Cre resulted in mouse lethality at postnatal (P) day 26.5^41^. Lamin-B2 expression stops after birth to promote maturation and early depletion of lamin-B2 causes defects in cardiomyocyte division late in embryonic development^42^. The role of lamin-B1 in cardiomyocytes remains unknown. Since lamin-A and -B1 show persistent expression in cardiomyocytes, we investigated the cardiomyocyte-specific roles of these two lamins in mice by generating cardiomyocyte-specific knockouts using Nkx2-5Cre.

Nkx2-5Cre begins expression in cardiac progenitors at approximately embryonic day (E)7.5^43^ and deletes the floxed *Lmna* and *Lmnb1* genes during early embryogenesis. Mice with homozygous Nkx2-5Cre and heterozygous for lamin-A and/or lamin-B1 (*Nkx2-5^Cre+/+^*;*Lmna^+/−^*;*Lmnb1^+/−^*or *Nkx2-5^Cre+/+^*;*Lmnb1^+/−^*) were crossed with mice homozygous for floxed lamin-A and/or lamin-B1 (*Lmna^f/f^*;*Lmnb1^f/f^*or *Lmnb1^f/f^*). These matings generated six cardiomyocyte-specific genotypes (Figure 1A): lamin-A and -B1 double heterozygotes (*Lmna^+/-^*; *Lmnb1^+/-^*), lamin-A heterozygote with lamin-B1 null (*Lmna^+/-^*;*Lmnb1^-/-^*), lamin-A null with lamin-B1 heterozygote (*Lmna^-/-^*;*Lmnb1^+/-^*), double null for lamin-A and lamin-B1 (*Lmna^-/-^*;*Lmnb1^-/-^*), lamin-B1 null (*Lmnb1^-/-^*) or lamin-B1 heterozygote (*Lmnb1^+/-^*). Deletion of either lamin-B1 or lamin-A floxed alleles reached an efficiency of >90% by E13.5, as determined by immunofluorescence staining for the relevant lamin proteins (Figure 1B–1E).

**Figure 1:**
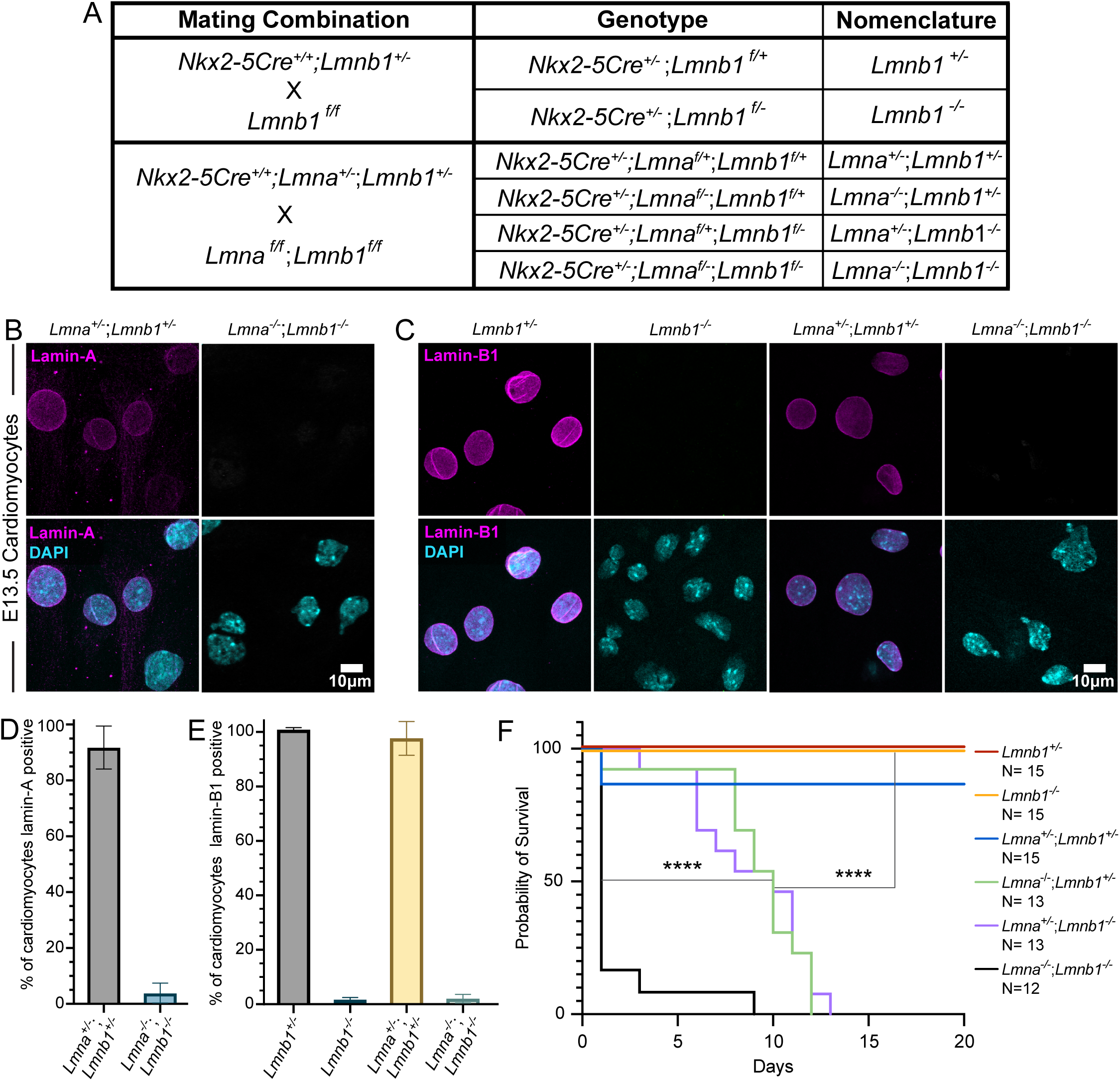
Lamin-A and lamin-B1 cooperatively regulate cardiomyocyte survival and postnatal viability. (A) Mating strategy to create lamin-A and -B1 genotypes. (B,C) Representative immunofluorescence images of cultured E13.5 cardiomyocytes stained for lamin-A (B) or lamin-B1 (C) (magenta) with DAPI (cyan), showing efficient Nkx2-5Cre-mediated depletion. Scale bar, 10 μm. (D,E) Quantification of lamin-A (D) and lamin-B1 (E) fluorescence intensity, demonstrating >90% depletion in knockout E13.5 cardiomyocytes (mean ± SD; n = 3 embryos per genotype). (F) Kaplan-Meier survival analysis of cardiomyocyte-specific lamin knockouts. Double knockout (*Lmna^-/-^*;*Lmnb1^-/-^*) mice exhibit perinatal lethality (∼P1), whereas mice retaining a single allele of lamin-A or lamin-B1 (*Lmna*^+/-^;*Lmnb1*^-/-^ or *Lmna*^-/-^;*Lmnb1*^+/-^) survive until P7-P14; *Lmnb1*^-/-^ mice are viable. Statistical significance was determined by log-rank (Mantel-Cox) test (****P < 0.0001).

We assessed mouse viability of each genotype using Kaplan-Meier survival analysis^44^ and found that most lamin-A and -B1 double knockout mice died at postnatal (P) day 1 (Figure 1F). The mice with only one allele of lamin-A (*Lmna^+/-^*;*Lmnb1^-/-^*) or lamin-B1 (*Lmna^-/-^*;*Lmnb1^+/-^*) survived until P7-14, while mice with lamin-B1 single knockout (*Lmnb1^-/-^*) were viable. Since lamin-A and -B1 double knockout mice die earlier than the other lamin-A and -B1 knockout combinations, the two lamins compensate for each other in cardiomyocytes to extend postnatal viability, with lamin-A playing a more important role than lamin-B1.

### Lamin-A and lamin-B1 are required for the organization of NPCs, LINC complexes, and chromatin

Previous studies have shown that the lamin meshwork counteracts forces generated by microtubule-based motors to ensure an even distribution of NPCs and LINC complexes around the nuclear envelope^18,19^. Visually, the E13.5 cardiomyocytes deleted of both lamin-A and -B1 (*Lmna^-/-^*;*Lmnb1^-/-^*) showed the strongest asymmetric distribution of NPCs and LINC complexes, followed by the lamin-B1 knockout with heterozygous lamin-A (*Lmna^+/-^*;*Lmnb1^-/-^*), and then the single lamin-B1 knockout (*Lmnb1^-/-^*) (Figure 2A, 2B). This is further confirmed by quantification (Figure 2C-E, see method). By contrast, cardiomyocytes containing lamin-A knockout with heterozygous lamin-B1 (*Lmna^-/-^*;*Lmnb1^+/-^*) showed even distributions of both NPCs and LINC complexes and were similar to controls (*Lmnb1^+/-^* and *Lmna^+/-^;Lmnb1^+/-^*) (Figure 2A, 2B, quantification not shown). These results could mean that lamin-B1 plays a more important role in maintaining an even distribution of NPCs and LINC complexes in E13.5 cardiomyocytes. However, Lamin-A is known to be expressed at a low level in embryonic cardiomyocytes^39^ and the total lamin amount, not a given lamin isoform, is required to ensure even distribution of NPCs in mouse embryonic fibroblasts (MEFs)^19^. Therefore, our findings suggest that the low amount of lamin-A in E13.5 cardiomyocytes renders these cells primarily reliant on lamin-B1 to maintain a crucial concentration of lamins required for an even distribution of NPCs and LINC complexes.

**Figure 2.**
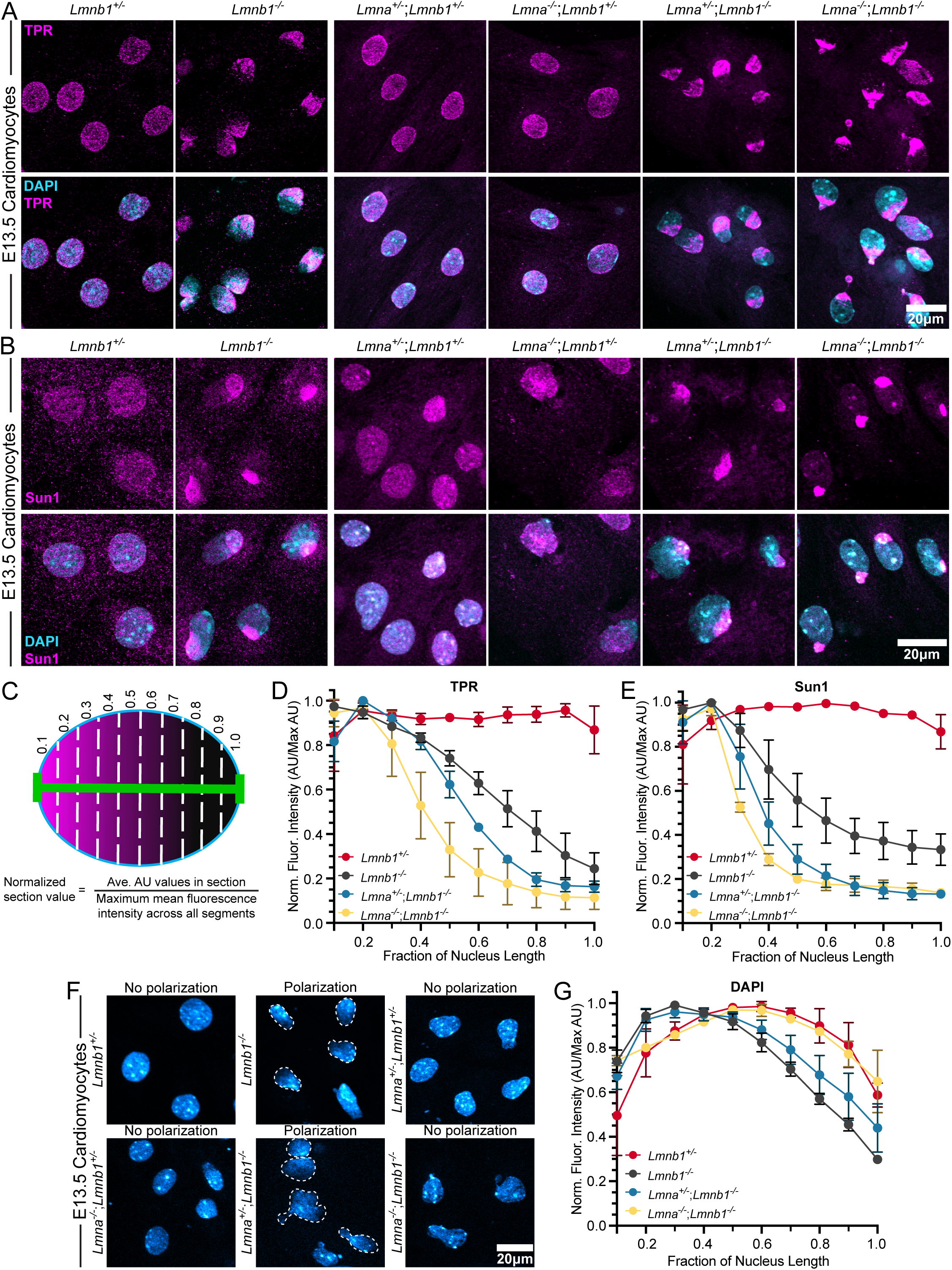
Lamin-A and lamin-B1 cooperate to maintain even distribution of NPCs, LINC complexes, and chromatin in E13.5 cardiomyocytes. (A, B) Representative confocal images of cultured E13.5 cardiomyocyte nuclei immunostained for either the NPC marker TPR (A, magenta) or the LINC complex component Sun1 (B, magenta) and counterstained with DAPI (cyan) across the indicated genotypes. Scale bar, 20 μm. (C) Schematic illustrating quantification of nuclear fluorescence distribution. Each nucleus was divided into ten equal sections along the nuclear axis (0.1–1.0), from highest to lowest signal when possible. Mean fluorescence was calculated per section and normalized to the maximum value within each nucleus, with 1.0 representing peak intensity. (D,E,G) Quantification of the normalized fluorescence intensity distribution for TPR (D), Sun1 (E), and DAPI (G). Mean ± SD; n = 3 biological replicates per genotype and stain (F) Representative confocal images of DAPI-stained nuclei in cultured E13.5 cardiomyocytes of the indicated genotypes. Scale bar, 20 μm.

Since lamin-A levels increase substantially in postnatal cardiomyocytes^39^, we examined whether this increase could rescue the distribution defects of NPCs and LINC complexes in genotypes that are lamin-B1 null with one or two copies of lamin-A. At P7, we found that distribution of NPCs and LINC complexes became even in *Lmnb1^-/-^* cardiomyocytes (Figure S1). However, cardiomyocytes with one copy of lamin-A and no lamin-B1 (*Lmna^+/-^*;*Lmnb1^-/-^*) continued to exhibit distribution defects in NPCs and LINC complexes (Figure S1). This shows that postnatal increase in lamin-A expression from both gene copies provides enough lamin protein to ensure proper distribution of NPCs and LINC complexes in the absence of lamin-B1. It further highlights the shared role of lamin-A and lamin-B1 in organizing the nuclear periphery in cardiomyocytes and emphasizes the importance of the total lamin concentration, not a specific lamin isoform, in ensuring proper distribution of NPCs and LINC complexes.

The polarization of NPCs and LINC complexes in the lamin-null MEFs were caused by dynein-mediated microtubule forces likely acting on these complexes and pulling them toward the minus ends of microtubules at the centrosome^18^. In embryonic cardiomyocytes, microtubules nucleated from centrosomes form a cage around the nucleus and radiate throughout the cytosol^45^. In postnatal cardiomyocytes, the microtubule cage surrounding the nucleus is maintained as the nucleus itself becomes the microtubule-organizing center. The microtubules radiating from the nucleus further reorganize into parallel arrays aligning with cardiac sarcomeres^46^. Most E13.5 cardiomyocytes have perinuclear microtubule cages and cytoplasmic microtubule arrays. However, we observed a small subpopulation of cells having centrosome-nucleated astral microtubule arrays with weak or no perinuclear microtubule cages and this was more abundant in certain Lamin-A and -B1 knockout combinations. The percentage of astral microtubule arrays in E13.5 cardiomyocytes, but no nuclear microtubule cage was greatest in cardiomyocytes without lamin-B1 (*Lmnb1^-/-^*, *Lmna^+/-^*;*Lmnb1^-/-^*and *Lmna^-/-^*;*Lmnb1^-/-^*) (Figure S2), and these microtubule asters appeared adjacent to the polarized NPCs and LINC complexes (Figure S2A, S2B). Notably, we found that the microtubule phenotype in lamin-B1 single knockout was rescued in postnatal cardiomyocytes, similar to the rescue of NPCs and LINC complexes (data not shown).

Since NPCs and LINC complexes are connected to chromatin at the nuclear periphery, their dramatic polarization could influence chromatin distribution^24^. We used DAPI staining to visualize cardiomyocyte DNA content in lamin-A and -B1 knockout combinations. *Lmnb1^-/-^* and *Lmna^+/-^*;*Lmnb1^-/-^* cardiomyocyte nuclei had noticeable polarization of DNA content, but surprisingly, this was not observed in the lamin-A and -B1 double knockout (*Lmna^-/-^*;*Lmnb1^-/-^*) cardiomyocytes, as confirmed by quantification (Figure 2F, 2G). The polarization of DAPI occurred to the same side as the polarized NPCs, LINC complexes, and microtubules (S2A, S2B). Therefore, normal DNA distribution in cardiomyocytes appears to not simply depend on total lamin amount or lamin type, but require additional factors, such as antagonistic forces and interactions among the two lamins, nuclear periphery proteins, and chromatin.

### Lamin-A and -B1 maintain cardiomyocyte nuclear shape with lamin-A required for preventing DNA damage in postnatal mouse cardiomyocytes

Lamin mutants have long been known to affect nuclear morphology, but few studies have examined this in the context of multiple lamin gene loss^24^. We examined the morphology of E13.5 cardiomyocyte nuclei from our different genotypes by measuring the overall roundness of the nuclei (see methods). Lamin knockout nuclei had generally greater disruption of nuclear morphology compared to controls (*Lmnb1^+/-^*and *Lmna^+/-^;Lmnb1^+/-^*). The lamin-A and -B1 double knockout (*Lmna^-/-^*;*Lmnb1^-/-^*) had the least round nuclei, followed by *Lmna^+/-^*;*Lmnb1^-/-^*, *Lmna^-/-^*;*Lmnb1^+/-^* and *Lmnb1^-/-^*knockouts (Figure 3A, 3B). Since mice with lamin-B1 cardiomyocyte knockout are fully viable, the presence of significant nuclear distortion is surprising, as similar distortion seen in *Lmna^-/-^*;*Lmnb1^-/-^*, *Lmna^-/-^*;*Lmnb1^+/-^*, and *Lmna^+/-^*;*Lmnb1^-/-^*cardiomyocytes coincided with mouse lethality (Figure 1F, 2F, 2G).

**Figure 3.**
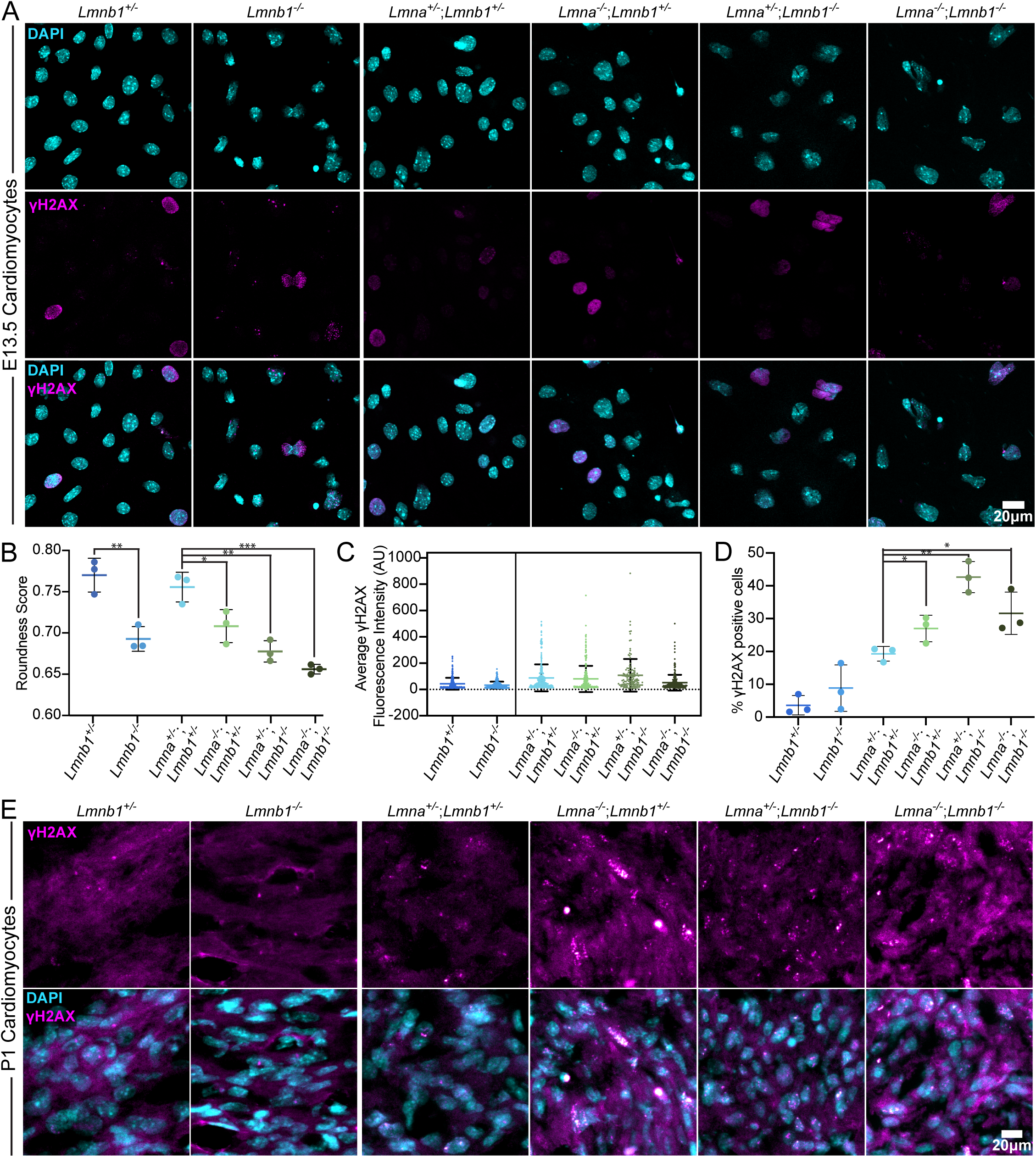
Lamin-A and lamin-B1 maintain cardiomyocyte nuclear shape, with lamin-A required for preventing DNA damage in postnatal cardiomyocytes. (A,E) Representative confocal images of cultured E13.5 (A) and P1 (E) cardiomyocyte nuclei immunostained for γH2AX (magenta) and counterstained with DAPI (cyan) across the indicated genotypes. Scale bar, 20 μm. (B) Mean and SD plot showing quantification of nuclear roundness scores in E13.5 cardiomyocytes across the indicated genotypes. Each dot represents one biological replicate with cells averaged per replicate. Mean ± SD; n = 3 biological replicates per genotype (Student’s t-test; *p < 0.05, **p < 0.01, ***p < 0.001). (C) Quantification of average γH2AX fluorescence intensity per nucleus in E13.5 cardiomyocytes across the indicated genotypes. Each dot represents one biological replicate with cells averaged per replicate. Mean ± SD; n = 3 biological replicates per genotype. No statistically significant differences were detected between genotypes (Student’s t-test). (D) Quantification of γH2AX positive cells per nucleus in P1 heart sections across the indicated genotypes. Mean ± SD; n = 3 biological replicates per genotype. (Student’s t-test; *p < 0.05, **p < 0.01).

Since lamin-A knockout in cardiomyocytes causes DNA damage^24,27^, we examined DNA damage in our knockout combinations by staining for double-stranded DNA breaks using γH2AX in E13.5 cardiomyocytes. We found that there was no significant increase in DNA damage in E13.5 cardiomyocytes in all four of our knockout combinations as compared to control single or double heterozygotes (Figure 3A, 3C). To assess whether DNA damage is associated with the onset of postnatal lethality, we analyzed cardiomyocytes of P1 mice. Staining with γH2AX revealed elevated DNA damage in *Lmna^-/-^*;*Lmnb1^+/-^, Lmna^+/-^*;*Lmnb1^-/-^*, and *Lmna^-/-^*;*Lmnb1^-/-^* cardiomyocytes compared to controls, while *Lmnb1^-/-^* cardiomyocytes were comparable to controls (Figure 3D, 3E). These findings indicate that lamin-A is required to prevent postnatal DNA damage in cardiomyocytes and that its deletion causes postnatal lethality in mice with *Lmna^-/-^*;*Lmnb1^+/-^*, *Lmna^+/-^*;*Lmnb1^-/-^*, and *Lmna^-/-^*;*Lmnb1^-/-^* genotypes.

### Lamin-B1 supports cardiomyocyte maturation and suppresses early embryonic gene expression programs

Although lamin-B1 knockout in cardiomyocytes is compatible with life (Figure 1F), the microtubule organization defects and DNA polarization (Figure 2F, 2G, S2), prompted us to explore whether lamin-B1 plays a role in the cardiomyocyte development. Since microtubules are required during development for establishing functionally important cardiomyocyte structures such as the cardiac dyad^46^, we used an antibody to Ryanodine Receptor 2 (Ryr2), a component of the dyads^47,48^ to analyze this structure. At E13.5, Ryr2-labeled cardiac dyad striations begin to form in control *Lmnb1^+/-^*cardiomyocytes as expected, but the *Lmnb1^-/-^*cardiomyocytes were largely devoid of striations (Figure 4A). Despite apparent rescue of microtubule organization and NPCs/LINC complex distributions in P7 *Lmnb1^-/-^* cardiomyocytes, the cardiac dyad striation remained disrupted with regions containing clustered and disorganized Ryr2 staining, while control *Lmnb1^+/-^* cardiomyocytes displayed a regular striated pattern (Figure 4B).

**Figure 4.**
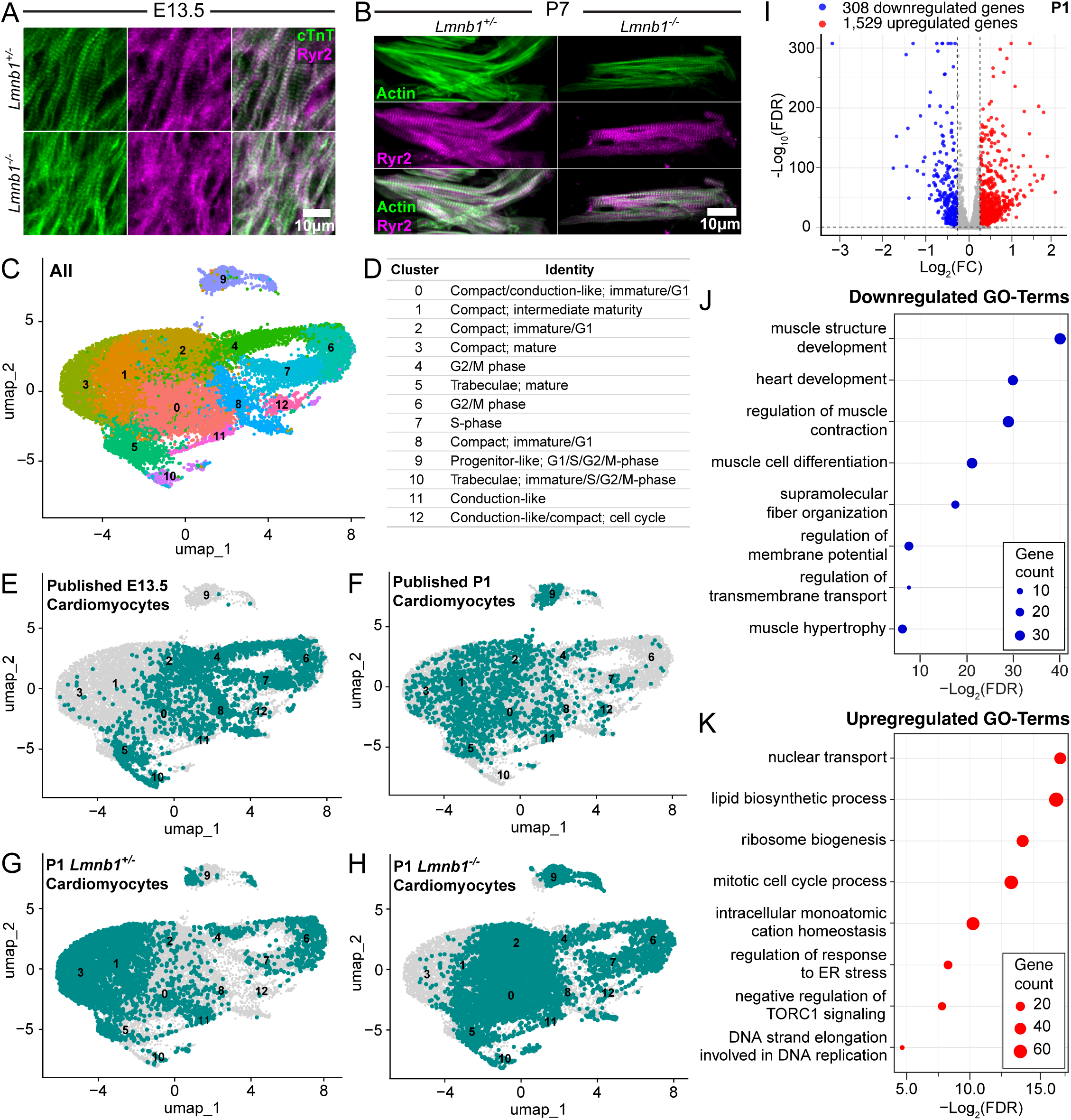
Lamin-B1 loss disrupts cardiac dyad organization and delays maturation and cell cycle exit of ventricular compact cardiomyocytes. (A,B) Representative confocal images of cultured E13.5 cardiomyocytes (A) and disrupted P7 tissue sections (B) immunostained for cardiac troponin T (cTnT, green) and Ryanodine Receptor 2 (Ryr2, magenta) in *Lmnb1^+/-^* and *Lmnb1^-/-^* cardiomyocytes. Scale bars are as indicated. N = 3 biological replicates per genotype and timepoint. (C) UMAP of all ventricular cardiomyocytes from the integrated previously published and P1 datasets, colored by cluster identity. (D) Table summarizing the identity of 13 ventricular cardiomyocyte clusters identified by scRNA-seq. (E-H) UMAPs showing the distribution of ventricular cardiomyocytes from the previously published E13.5 (E) and P1 (F) datasets and P1 *Lmnb1^+/-^*(G) and *Lmnb1^-/-^* (H) datasets (teal), projected onto the integrated UMAP combining previously published datasets and *Lmnb1^+/-^*and *Lmnb1^-/-^* datasets, with all other cells shown in gray. (I) Volcano plot of differentially expressed genes in compact ventricular cardiomyocytes from *Lmnb1^-/-^*versus *Lmnb1^+/-^* hearts. Downregulated genes and upregulated genes are shown in blue and red, respectively. The dashed lines indicate significance thresholds (|Log_2_FC| > 0.26; p < 0.05). Differential gene expression analysis was restricted to genes expressed in more than 25% of cells in at least one genotype. (J, K) Selected GO-terms enriched among downregulated (J) and upregulated (K) genes in *Lmnb1^-/-^* compact ventricular cardiomyocytes.

Since *Lmnb1^-/-^* cardiomyocytes do not exhibit DNA damage, we have an opportunity to study how lamin-B1 can influence gene expression without the confounding effect of DNA damage. We first performed bulk RNA-seq on whole P7 hearts and found 280 upregulated and 198 downregulated genes in *Lmnb1^-/-^* hearts compared to controls (Figure S3A, Table 1). Interestingly, the gene ontology (GO) analyses show that the downregulated genes were enriched for cardiomyocyte-specific processes, including contraction, sarcomere assembly, and cardiac dyad organization (Figure S3B, Table 1), which are consistent with impaired maturation^36,49^. The upregulated genes were dominated by those associated with cell cycle and early metabolic processes (Figure S3C, Table 1). This suggests that lamin-B1 plays a role in the timely expression or repression of genes involved in the maturation and intracellular organization of cardiomyocytes.

Next, we used single cell RNA-sequencing (scRNA-seq) of P1 hearts to analyze cardiomyocyte-specific effects of lamin-B1 knockout. P1 hearts were selected because the cardiomyocytes are known to begin the maturation process at this time point^36^. We integrated our dataset with published scRNA-seq datasets from nine wild type embryonic time points (E10.5-E18.5) and eight postnatal timepoints (P1-P6, P8 and P9)^50,51^, which helped us to define cell types and assess the developmental stages of our lamin-B1 knockout and control cardiomyocytes. Initial integration resolved major heart cell populations, including cardiomyocytes, epicardial cells, endothelial cells, fibroblasts, and immune cells (Figure S4A). We then focused on cardiomyocytes based on their expression of titin (*Ttn*+) and alpha-actinin-2 (*Actn2*+)^51^. The atrial and ventricular cardiomyocytes are transcriptionally divergent^50–52^. Since ventricular cardiomyocytes represent the major population of cardiomyocytes in the heart^52^, which is also reflected in our datasets, we focused on ventricular cardiomyocytes for our analyses. Based on their expression of myosin light chain 2 (*Myl2+*) and the lack of expression for sarcolipin (*Sln-*)^51^, we identified 4,307 *Lmnb1^+/-^* and 6,641 *Lmnb1^-/-^* ventricular cardiomyocytes.

During heart ventricular development, the cardiomyocyte progenitors differentiate into trabecular and compact myocardial populations, which form the inner trabeculated layer and the outer compact ventricular wall, respectively. In addition, a smaller subset of cardiomyocytes exhibits conduction-like transcriptional features and these cells contribute to the synchronization of cardiac contraction^52,53^. Re-clustering of ventricular cardiomyocytes identified 13 transcriptionally distinct clusters corresponding to compact, trabecular, and conduction-like populations (Figure 4C, 4D). Cell identities were assigned using known marker expression and GO-term enrichment analysis^50,51,54^, while the cell-cycle phase was determined using Seurat-based cell-cycle scoring. The compact cardiomyocytes were represented by clusters 0, 1, 2, 3, and 8; trabecular cardiomyocytes by clusters 5 and 10; conduction-like populations by clusters 11 and 12; and a progenitor-like state by cluster 9. In addition to compact features, Clusters 0 and 8 displayed conduction-associated features (Figure 4C, 4D, S4B). The compact cardiomyocytes in clusters 4, 6, and 7 had expression profiles indicating they are in S-phase (cluster 7) or G2/M-phase (clusters 4 and 6) of the cell cycle (Figure 4C, 4D, S4C). We focused our subsequent analyses on the compact cardiomyocytes as they make up the majority of cardiomyocytes and undergo significant maturation and remodeling starting from P1.

The published scRNA-seq datasets cover a broad developmental time window^50,51^, which facilitates the developmental staging of compact cardiomyocytes from our datasets. Clusters 2 and 8 represent immature compact cardiomyocytes, marked by active cell cycle progression and transcriptional repression of maturation (Figure 4C, 4D, S4B). Cluster 0 represents a distinct compact population with conduction-like identity, characterized by conduction system specification and ECM remodeling genes (Figure 4C, 4D, S4B). Cluster 1 exhibits an intermediate maturation state marked by upregulation of fatty acid oxidation and metabolic reprogramming (Figure 4C, 4D, S4B). Cluster 3 represents the most mature compact cardiomyocyte population, with high expression of genes associated with sarcomeric maturation and hypertrophic growth (Figure 4C, 4D, S4B). The compact cardiomyocytes undergoing cell cycle are found in clusters 4, 6, and 7 (Figure 4C, 4D, S4C). Consistent with these classifications, the published wild type E13.5 compact cardiomyocytes are enriched in the immature (clusters 0, 2, 8) and cell cycle clusters (clusters 4, 6, 7) (Figure 4E). The published postnatal P1 compact cardiomyocytes were enriched in intermediate (cluster 1) and mature (cluster 3) clusters, consistent with postnatal cardiomyocyte maturation and cell-cycle exit (Figure 4F). We overlayed our scRNA-seq of P1 compact cardiomyocytes from our control and lamin-B1 knockout mice onto the published datasets. As expected, we found compact cardiomyocytes from our lamin-B1 control were distributed across all clusters but were particularly enriched in cluster 1 and cluster 3 that contain more mature cardiomyocytes and cardiomyocytes within an intermediate maturation stage, respectively (Figure 4G, S4D). By contrast, lamin-B1 knockout cardiomyocytes showed a depletion of cluster 3 and a reduction in cluster 1, but instead were enriched in immature clusters 0, 2, and 8 (Figure 4H, S4D). This shows that the lamin-B1 cardiomyocyte-specific knockout resulted in delayed maturation and cell cycle exit of compact cardiomyocytes.

To examine how lamin-B1 knockout impacted transcription in ventricular compact cardiomyocytes, we combined clusters 0-4 with clusters 6-8 and analyzed differential gene expression upon lamin-B1 knockout (Figure 4I, Table 2). GO-term analysis reveal that the downregulated genes are involved in cardiomyocyte maturation, including those encoding proteins found in sarcomere and cardiac dyad structures (Figure 4J, Table 2), while the upregulated genes are involved in cell cycle, embryonic metabolic processes, ion handling, and negative regulation of hypertrophy (Figure 4K, Table 2). Therefore, loss of lamin-B1 results in a delay in maturation of compact cardiomyocytes characterized by prolonged expression of cell cycle genes and the downregulation of genes involved in cardiomyocyte structural maturation and contraction.

### Lamin-B1 maintains LADs and chromosome territories in cardiomyocytes

We showed that lamins organize LADs, which in turn affects 3D chromatin interactions throughout the genome^55^. To understand how lamin-B1 could support expression of genes involved in cardiomyocyte maturation, we first explored whether its deletion disrupted cardiomyocyte LADs. We mapped LADs using Cleavage Under Targets and Release Using Nuclease (CUT&RUN) with antibodies targeting lamin-A in control and lamin-B1 knockout P1 cardiomyocytes (see methods). LAD and non-LAD regions were then defined using a two-state hidden Markov model (HMM) (Figure 5A). Upon lamin-B1 loss, chromosomes exhibited an average ∼23% reduction in LAD coverage across the cardiomyocyte genome (Figure 5B). The short LAD regions (median length ∼155 kb) tended to show complete detachment from the nuclear lamina, whereas the long LAD regions (median length ∼975 kb) showed partial detachment primarily at their edges (Figure 5A, 5C). Thus, lamin-B1 is required for maintaining proper LADs organization in cardiomyocytes.

**Figure 5.**
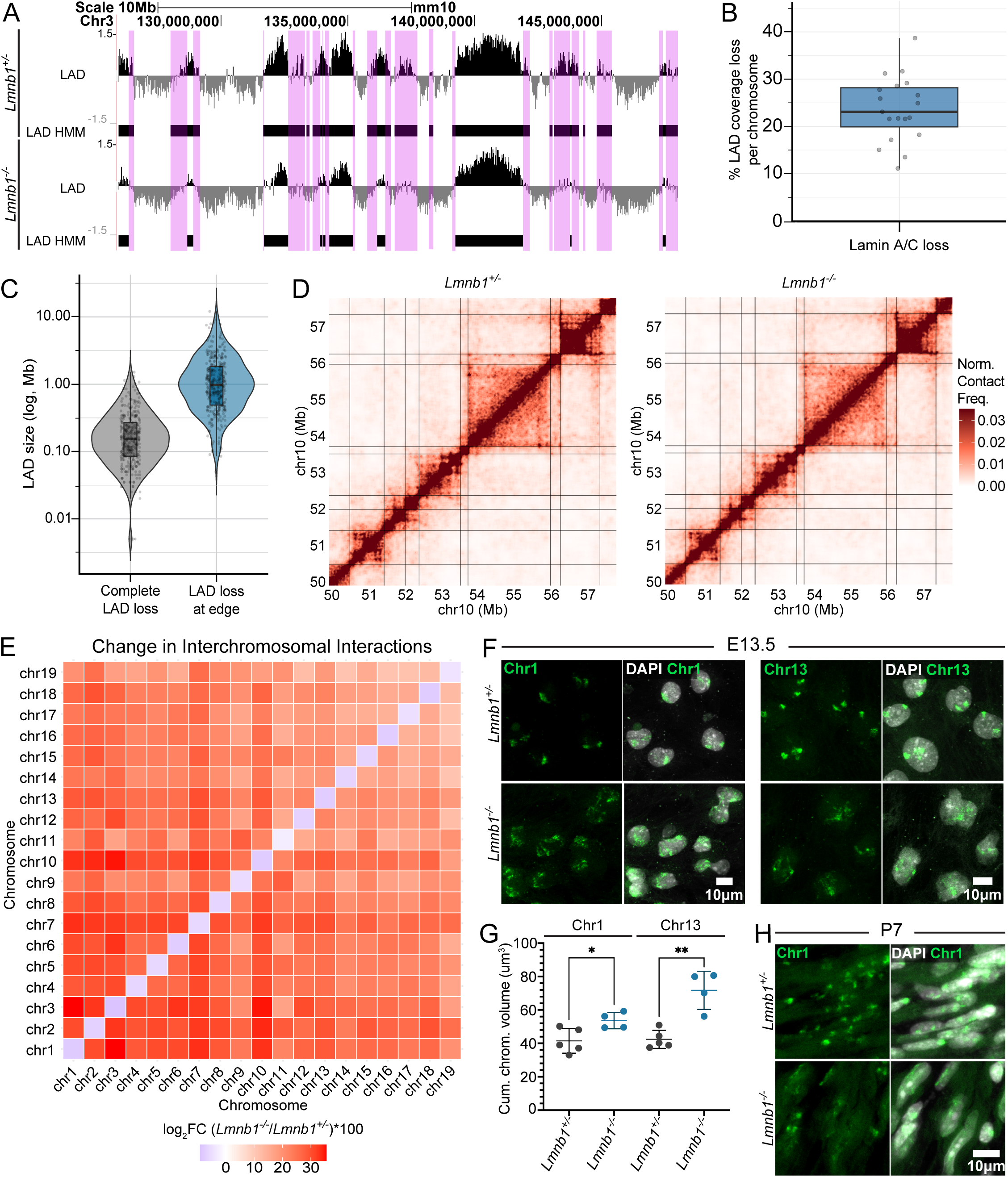
Lamin-B1 maintains LADs and chromosome territories in cardiomyocytes. (A) Genome browser tracks showing LADs mapping with lamin-A CUT&RUN z-score signal and HMM-defined LAD calls (black bars) for a representative region of chromosome 3 in *Lmnb1^+/-^* (top) and *Lmnb1^-/-^*(bottom) P1 cardiomyocytes. Example regions of LAD loss are highlighted in purple blocks. (B) Box and whisker plot showing the percent loss of LAD coverage across each chromosome between *Lmnb1^-/-^*and *Lmnb1^+/-^* cardiomyocytes. Gray dots represent individual chromosomes. (C) Violin plots showing the size distribution (log scale) of LADs undergoing complete loss versus loss at edges in *Lmnb1^-/-^* cardiomyocytes. Box plots within violins indicate median and interquartile range. (D) ICE-normalized Hi-C contact maps for a representative region of chromosome 10 in *Lmnb1^+/-^* (left) and *Lmnb1^-/-^* (right) P1 cardiomyocytes at 10 kb resolution. Color scale represents normalized contact frequency. (E) Interchromosomal interaction matrix showing the percent change in inter-chromosomal contacts in *Lmnb1^-/-^*relative to *Lmnb1^+/-^* cardiomyocytes. (F) Representative fluorescence in situ hybridization (FISH) images of chromosome 1 (Chr1, green, left) and chromosome 13 (Chr13, green, right) counterstained with DAPI (gray) in *Lmnb1^+/-^*and *Lmnb1^-/-^* cardiomyocytes at E13.5. Scale bar, 10 μm. (G) Quantification of cumulative chromosome volume (µm^3^) for Chr1 and Chr13 in *Lmnb1^+/-^*and *Lmnb1^-/-^*Quantification of cumulative chromosome volume (µm3) for Chr1 and Chr13 in *Lmnb1^+/-^*and *Lmnb1^-/-^* E13.5 cardiomyocytes. Each dot represents one biological replicate with cells averaged per replicate. Mean ± SD; n ≥ 4 biological replicates per genotype (Student’s t-test; *p < 0.05, **p < 0.01). (H) Representative FISH images of Chr1 (green) counterstained with DAPI (gray) in *Lmnb1^+/-^* and *Lmnb1^-/-^* P7 cardiomyocytes. Scale bar, 10 μm.

Next, we mapped 3D chromatin interactions by high-throughput chromosome conformation capture (Hi-C) in isolated P1 cardiomyocytes using our recently developed Hi-C method for small numbers of cells. We performed Hi-C in two biological replicates in lamin-B1 knockout and heterozygote littermates. After filtering out duplicates, invalid pairs, and unmapped reads, we obtained 147,005,252 reads for lamin-B1 heterozygote and 188,511,585 reads for lamin-B1 knockout cardiomyocytes (Table 3). All samples had >70% intrachromosomal contacts, indicating low noise from random ligation^56^ (Table 3). We generated normalized contact maps by applying the iterative correction and eigenvector decomposition (ICE) method^57^ (Figure 5D). To evaluate our Hi-C reproducibility, we assessed similarities at different resolutions and found high correlation between replicates (Figure S5A). We further plotted the contact frequency as a function of genomic distance, which showed a high consistency among all Hi-C replicates (Figure S5B). This shows a high reproducibility and quality of our Hi-C map despite low-cell input. Using insulation scores derived from ICE-normalized contacts, we identified TAD boundaries and found that >87% TADs were preserved between control and lamin-B1 knockout cardiomyocytes, revealing that TADs are mostly unaffected by lamin-B1 loss (Figure 5D, S5C, S5D).

We then performed compartment analyses using CscoreTool to assign scores and divide the cardiomyocyte genome into A and B compartments based on higher interactions within each compartment than between the two^58^. The A and B compartments correspond to genomic regions found in the interior and nuclear periphery, respectively^3–5,55^. The positive C-scores represent regions within the B compartment while the negative scores represent chromatin in the A compartment, and higher scores indicate stronger compartmentalization. Since LADs showed high correlation with the B compartment as expected (Figure S5E, S5F, Pearson value = 0.73), we compared compartment scores between lamin-B1 knockout and control cardiomyocytes. Consistent with the observed LAD loss, we found that lamin-B1 knockout caused significant genome-wide shift of chromosome regions from compartment B towards A (Figure S5G) with the shift being most pronounced in regions that showed LAD loss upon lamin-B1 deletion (Figure S5H).

Individual chromosomes are folded to occupy spatially distinct chromosome territories within the nucleus. This is reflected by the substantially greater intrachromosomal than interchromosomal interactions^4,59^. Strikingly, we found that lamin-B1 loss caused a strong increase in inter-chromosomal interactions with larger chromosomes showing a greater increase than smaller chromosomes (Figure 5E). One interpretation is that lamin-B1 loss causes the unfolding of individual chromosomes, which could allow more inter-chromosomal interactions. To explore this, we performed fluorescent in situ hybridization (FISH) on whole chromosomes. We utilized chromosome paint probe mixes previously designed in our lab that target chromosomes 1 and 13^55^, representing the largest and a smaller chromosome, respectively. We measured chromosome volumes based on FISH signal using the Imaris machine learning segmentation tool. The tool was trained on each chromosome individually using images from both control and lamin-B1 knockout cardiomyocytes to accurately distinguish true signals over background. We first determined chromosome volumes in E13.5 cardiomyocytes and found noticeable expansion of chromosome territories in lamin-B1 knockout E13.5 cardiomyocytes as compared to the controls. This expansion often manifested as elongation of chromosomes across the length of the nucleus (Figure 5F, 5G). This chromosome expansion persisted in P7 cardiomyocytes (Figure 5H). Therefore, cardiomyocyte-specific lamin-B1 knockout results in a loss of a subset of LADs and marked chromosome expansion. The chromosome expansion persists from embryonic stages to at least the first week of postnatal development even though NPC and LINC complex distributions are fully rescued.

### Lamin-B1 can support cardiomyocyte gene expression by maintaining 3D chromatin interactions

Expression of a given gene is known to be influenced by its immediate chromatin neighborhoods. Neighborhoods that contain active epigenetic modifications and accessible chromatin support gene expression, while those containing repressive epigenetic modifications and condensed chromatin can repress gene expression^55,60^. To understand how lamin-B1 loss causes gene dysregulation relevant to cardiomyocyte function, we explored whether lamin-B1 maintains chromatin neighborhoods. We previously developed a <u>Hi</u>stone and lamina <u>Lands</u>cape (HiLands) model that classifies chromatin regions based on their epigenome and nuclear lamina association^61^, which can help infer whether a chromatin region is in an active or inactive chromatin neighborhood. We defined HiLands in control P1 cardiomyocytes utilizing previously published datasets for Histone H3 lysine trimethylation (H3K27me3), H3K4me1, and H3K4me3 mapped in wild type P1 cardiomyocytes^49^. We additionally performed CUT&RUN to map H3K9me2, H3K9me3 and H3K27ac in wild type P1 cardiomyocytes and ATAC-seq (Assay for Transposase-Accessible Chromatin sequencing) for chromatin accessibility in our control lamin-B1 heterozygous P1 cardiomyocytes. Using these datasets and our LADs mapping, we performed Hidden Markov Modeling (HMM) and identified 6 states as HiLands-Purple (P), - Blue (B), -Green (G), -Yellow (Y), -Orange (O), and -Red (R) (Figure 6A, 6B, S6A).

**Figure 6.**
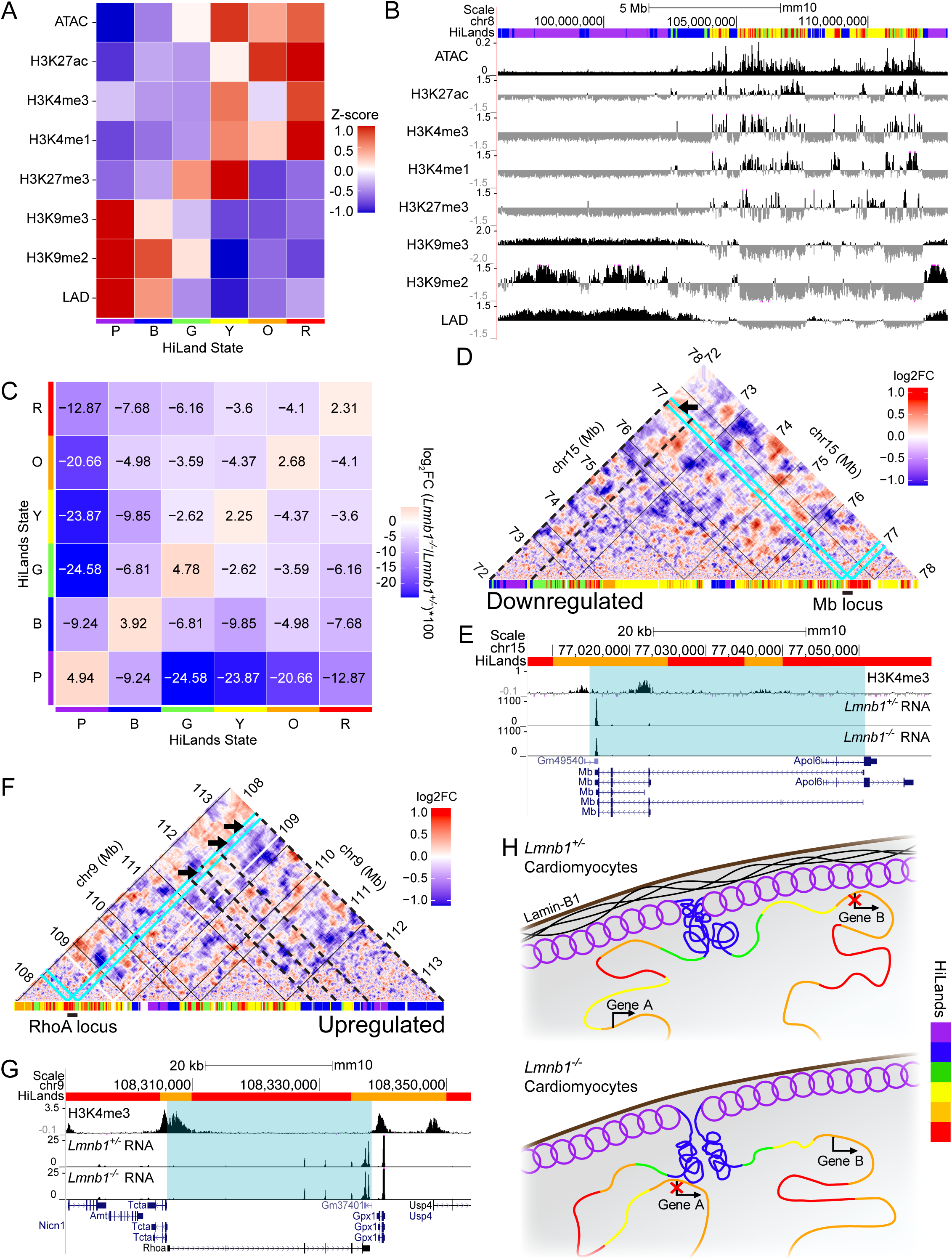
Lamin-B1 maintains chromatin neighborhoods that support cardiomyocyte gene expression. (A) Heatmap showing the mean z-score of lamin-A CUT&RUN LADs mapping, ATAC-seq, and different epigenomic modifications across the six HiLands states defined in wild-type and control *Lmnb1^+/-^* P1 cardiomyocytes. (B) Genome browser tracks showing LADs mapping, ATAC-seq and histone modification signals alongside a representative region of chromosome 8 in wild-type and *Lmnb1^+/-^* P1 cardiomyocytes. HiLands state annotation is shown at the top. (C) Matrix showing the change in chromatin interactions between pairs of HiLands states in *Lmnb1^-/-^* relative to *Lmnb1^+/-^* P1 cardiomyocytes. Values represent Log_2_FC × 100. (D,F) Hi-C contact maps (10 kb resolution, dynamic binning; see Methods) showing Log_2_ fold changes in chromatin interactions in *Lmnb1^-/-^*relative to *Lmnb1*^+/-^ cardiomyocytes at the Mb locus on chromosome 15 (D) and the RhoA locus on chromosome 9 (F). HiLands annotations are shown along the diagonal, with loci indicated by cyan lines and TAD boundaries indicated by solid black lines. Regions showing increased contact changes with HiLands-B/-P (D) or decreased contact changes with HiLand-B (F) are shown with dashed black lines and black arrows. (E,G) Genome browser tracks at the Hb locus (E) and RhoA locus (G) showing H3K4me3 modification, CPM-normalized scRNA-seq signal from *Lmnb1^+/-^*and *Lmnb1^-/-^* compact ventricular cardiomyocytes, and gene annotation. The HiLands states are shown at the top and the gene region is highlighted in cyan. (H) Schematic model of chromatin neighborhood disruption following lamin-B1 loss. In *Lmnb1^+/-^* cardiomyocytes (left), Gene A associates with active HiLands states and is expressed, while Gene B associates with repressive HiLands states and is silenced. In *Lmnb1^-/-^*cardiomyocytes (right), LAD detachment and chromosome expansion alter chromatin neighborhoods, causing Gene A to gain repressive contacts and become silenced, while Gene B loses repressive contacts and becomes expressed.

HiLands-B and -P corresponded to LADs and were enriched in H3K9me2, with HiLands-P also enriched for H3K9me3 (Figure 6A, 6B, S6B). As expected, gene expression, gene density, and chromatin accessibility as revealed by ATAC-seq were low in HiLands-B and -P LADs (Figure 6A, S6C-F). Interestingly, HiLands-G chromatin regions showed low levels of epigenetic modifications except for H3K27me3 followed by H3K9me2 but had elevated chromatin accessibility and gene density and lower C-score values compared to HiLands-P and -B (Figure 6A, S6B-G). This is consistent with HiLands-G chromatin being distributed mostly in the non-LAD regions but having low gene density and gene expression in cardiomyocytes (Figure S6B, S6F). HiLands-Y represent poised chromatin regions that are enriched in both H3K27me3 and H3K4me1 (or H3K4me3) (Figure 6A)^62^. HiLands-O and -R are enriched for H3K4me1 and H3K27ac, with HiLand-R also enriched for H3K4me3, a classic marker for active promoters (Figure 6A)^62^. HiLands-Y, -O, and -R all contain dense ATAC-seq peaks and the lowest C-score values, which is consistent with their interior chromatin features and high gene densities (Figure 6A, S6C-E, S6G). As expected, mapping of our P1 scRNA-seq from the control compact cardiomyocytes showed that HiLands-R and -O are enriched for expressed genes followed by HiLand-Y (Figure S6F).

To understand how lamin-B1 loss altered chromatin neighborhoods, we analyzed chromatin interaction changes based on HiLands. Upon lamin-B1 loss, we found decreased interactions between HiLands-P and HiLands-B (Figure 6C). In addition, HiLands-B showed a loss while HiLands-P showed a gain of LAD signal (Figure S6H), suggesting detachment of HiLands-B from the nuclear lamina. This coincided with decreased C-score values for HiLands-B than for HiLands-P, indicating a detachment of HiLands-B from the nuclear lamina (Figure 6I). This detachment may cause an interior shift of HiLands-G, -Y, -O, and -R, which can explain the overall significant reduction of interactions of these HiLands with HiLand-P and -B upon lamin-B1 loss (Figure 6C).

To explore whether chromatin interactions among HiLands may contribute to the transcriptional dysregulation seen in lamin-B1 null cardiomyocytes, we analyzed the interactions of promoters of dysregulated genes with their flanking HiLands states (see methods). We noticed that for both downregulated and upregulated genes, there was a significant tendency for their promoters to show decreased interactions with HiLands-B and -P. This could be caused by the overall shifting of interior chromatin regions represented by HiLands-G, -Y, -O, and -R away from HiLands-B and -P (Figure 6C). Upon comparing to non-dysregulated gene promoters, we found a significantly greater trend for downregulated gene promoters to have increased interactions with HiLand-B and -P (Figure S6J). This was accompanied by decreased interactions with HiLands-Y compared to non-dysregulated genes (Figure S6J), but these decreased interactions do not explain gene expression changes upon lamin-B1 loss. Since HiLands-B and -P are strongly associated with heterochromatin states, we looked at how many downregulated genes may be explained by these changes in interactions. Approximately 26.9% of downregulated genes had at least 1.2-fold increase of interactions with either HiLands-B or -P, suggesting that a subset of downregulated genes may be repressed through increased association with repressive chromatin neighborhoods (Table 4). An example of this gene is the gene *Mb*, which encodes myoglobin, a protein responsible for oxygen storage and transport that sharply increases after birth to support oxygen needs (Figure 6D, 6E)^63^. The remaining downregulated genes may be silenced through indirect mechanisms, such as downstream transcriptional cascades initiated by the chromatin neighborhood changes.

Conversely, for upregulated genes, we found their promoters showed significantly reduced interactions with HiLands-B and -O compared to non-dysregulated genes (Figure S6J). While changes in interactions with Hilands-O could not explain the upregulated genes, the reduced interactions with the repressive HiLands-B may explain a subset of the upregulated genes. We found approximately 29.2% of upregulated genes showed at least 1.2-fold decreased interactions with HiLands-B (Table 4). One such example was the gene *Rhoa*, which promotes differentiation and proliferation in the embryonic heart (Figure 6F, 6G)^64^. The reduced interactions with repressive HiLands-B chromatin neighborhoods can lead to gene upregulation. Some of these upregulated genes could lead to upregulation of additional downstream genes. Therefore, lamin-B1 loss may disrupt chromatin neighborhoods, leading to gene dysregulation in cardiomyocytes (Figure 6H).

### Chromatin neighborhoods maintained by lamin-B1 can influence transcription factors for cardiomyocyte maturation and cell cycle exit

We showed that loss of lamin-B1 resulted in prolonged expression of cell cycle genes and downregulation of genes involved in cardiomyocyte structural maturation and contraction (Figure 4). Transcription factor activities are known to be impacted by chromatin neighborhoods^65,66^. To explore how lamin-B1 maintenance of chromatin neighborhoods could in turn regulate gene expression in cardiomyocytes, we set out to identify binding motifs for transcription factors among the dysregulated genes upon lamin-B1 loss. We first used the previously published H3K4me3 datasets for wild type P1 compact cardiomyocytes to identify promoters for the dysregulated genes^49^. We then used HOMER (Hypergeometric Optimization of Motif EnRichment) motif finder to identify enriched binding motifs for transcription factors (Table 5). We focused on the top five most significantly enriched binding motifs for the up- and down-regulated genes (Figure 7A, 7B).

**Figure 7.**
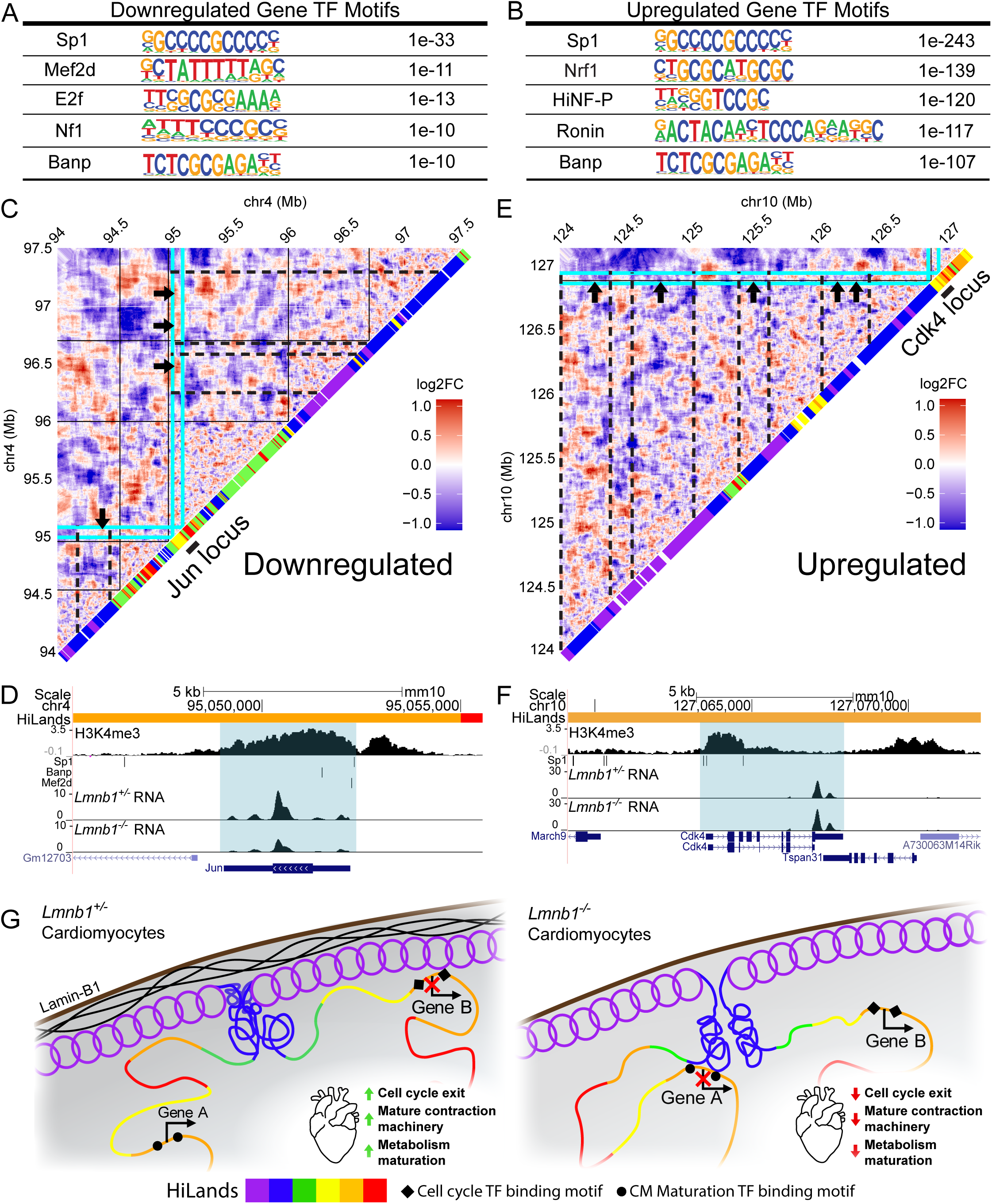
Lamin-B1 loss alters transcription factor binding environments at dysregulated gene loci. (A,B) The top five enriched transcription factor binding motifs identified by HOMER analysis at promoters of downregulated (A) and upregulated (B) genes in *Lmnb1^-/-^* compact ventricular cardiomyocytes. (C, E) Hi-C contact maps (10 kb resolution, dynamic binning; see Methods) showing Log_2_ fold changes in chromatin interactions in *Lmnb1^-/-^*relative to *Lmnb1*^+/-^ cardiomyocytes at the Jun locus on chromosome 4 (C) and the Cdk4 locus on chromosome 10 (E). HiLands annotations are shown along the diagonal, with loci indicated by cyan lines. HiLands annotations are shown along the diagonal, with loci indicated by cyan lines and TAD boundaries indicated by black lines. Regions showing increased contact changes with HiLands-B/-P (C) or decreased contact changes with HiLand-B (E) are shown with dashed black lines and black arrows. (D, F) Genome browser tracks at the Jun locus (D) and Cdk4 locus (F) showing H3K4me3 modification, binding motif positions for Sp1 (Jun and Cdk4), Banp (Jun), and Mef2d (Jun), CPM-normalized scRNA-seq signal from *Lmnb1^+/-^*and *Lmnb1^-/-^* compact ventricular cardiomyocytes, and gene annotation. The HiLands states are shown at the top and the gene region is highlighted in cyan. (G) Schematic model illustrating how lamin-B1 loss alters transcription factor binding environments. In *Lmnb1^+/-^* cardiomyocytes (left), cardiomyocyte maturation transcription factor binding motifs (black circles) associate with active chromatin regions, while cell cycle/early developmental motifs (black diamonds) associate with repressive chromatin regions, promoting cardiomyocyte maturation. In *Lmnb1^-/-^* cardiomyocytes (right), chromatin neighborhood shifts reduce maturation-associated motif availability to transcription factors and increase access to motifs linked to cell cycle and immature gene programs, impairing maturation.

Interestingly, both the up- and down-regulated genes are significantly enriched for binding motifs for transcription factor Sp1 and Banp. Sp1 has established roles in regulating early cardiomyocyte differentiation, proliferation, and later metabolic shift^67^. Meanwhile, Banp (also called Smar1) has been shown to influence cell division and prevent cell death^68,69^. The down-regulated genes are further enriched for binding motifs associated with transcription factors known to regulate cell cycle (E2f^70^ and Nf1^71^), and cardiomyocyte cell cycle exit and apoptosis prevention (Mef2d)^72^ during development. In addition to Sp1 and Banp, the up-regulated genes are also significantly enriched in binding motifs for transcription factors involved in midgestational cardiogenesis (Ronin)^73^, general control of G1/S phase transition during rapid embryonic cell divisions (HiNF-P)^74^, and neonatal heart regeneration (Nrf1)^75^.

For downregulated genes, 17.5% had significantly increased interactions with HiLand-B and -P and contain at least one of the top five binding motifs (Figure 7A, Table 5). Meanwhile, 20.5% of all upregulated genes that have at minimum one of the top five binding motifs exhibit significantly decreased interactions with Hiland-B (Figure 7B, Table 5). This suggests that lamin-B1 loss can bring genes into a transcriptionally repressive chromatin neighborhood, which can impact the binding or activities of the transcription factors to their motifs on the promoters of these genes, leading to their downregulation. For example, lamin-B1 loss caused increased interactions between the promoter of the gene *Jun* with HiLands-B and -P (Figure 7C). This may reduce the binding or activities of transcription factors Sp1, Banp, and Mef2d that regulate *Jun*, leading to *Jun* downregulation (Figure 7D). Conversely, lamin-B1 loss caused a decreased interaction between the promoter for the gene *Cdk4* (Figure 7E) with HiLands-B. This could increase the binding or activities of the transcription factor Sp1 to *Cdk4*, resulting in *Cdk4* upregulation (Figure 7F).

We next analyzed whether dysregulation of *Jun* and *Cdk4* could explain other dysregulated genes that do not show chromatin neighborhood change upon lamin-B1 loss. We found Jun binding motifs were present in 16.2% of these genes (Table 4). Therefore, the downregulation of *Jun* expression due to chromatin neighborhood changes could in turn lead to down regulation of these genes. Since Jun is known to promote cardiomyocyte maturation in postnatal hearts^49^, its reduction could lead to the downregulation of additional genes required for cardiomyocyte maturation.

For the up-regulated genes that cannot be explained by chromatin neighborhood changes due to lamin-B1 loss, the increase in Cdk4 could provide an explanation. Cdk4 is known to phosphorylate Rb, leading to the release of E2F transcription factors. Among the E2Fs, E2F3 is known to promote cardiomyocyte proliferation^70,76^. Thus, releasing E2F proteins from Rb could lead to upregulation of E2F target genes. We found that 42.5% of the upregulated genes that do not show chromatin neighborhood changes contain E2F3 binding motifs (Table 4). Therefore, upregulation of Cdk4 may support the upregulation of these genes.

In total, of all the dysregulated genes caused by lamin-B1 loss in cardiomyocytes, 38.0% can be explained by chromatin neighborhood change-related alteration of transcription factor activities. An additional 58.7% can be explained by the dysregulation of the binding or activities of Jun, Cdk4, and E2F3. Therefore, our findings support a mechanistic framework linking lamin disruption with chromatin neighborhood reorganization, which affects transcription factor activity and transcriptional programs that promote cardiomyocyte structural maturation and cell cycle exit (Figure 7G).

## Discussion

Despite the importance of lamins in heart development, our knowledge is limited to the role of either lamin-A or -B2 in cardiomyocytes. Neither the cardiomyocyte-specific role of lamin-B1 nor the question of whether the different lamin isoforms have compensatory functions have been addressed. By dissecting an allelic series of cardiomyocyte-specific knockouts of lamin-A and lamin-B1, we resolve both questions and reveal a distinctive role for lamin-B1 in coupling 3D genome organization to the transcriptional logic of cardiomyocyte maturation and cell-cycle exit. Lamin-B2 was excluded from our allelic analysis because its expression is rapidly silenced shortly after birth, the time in which cardiomyocytes undergo terminal maturation and cell-cycle exit. Nonetheless, our findings suggest that the two B-type lamins have evolved different roles with lamin-B1 supporting cell cycle exit and lamin-B2 promoting cell division.

Our analyses of the allelic series reveal compensatory functions between lamin-A and - B1. The double knockout (*Lmna^-/-^*;*Lmnb1^-/-^*) mice die at ∼P1, while either single homozygote with one remaining wild-type allele of the other lamin survive an additional 1-2 weeks and lamin-B1 single knockout (*Lmnb1^-/-^*) mice are viable (Figure 1F). At E13.5, when lamin-A levels in cardiomyocytes are low, the distributions of NPCs and LINC complexes in the double knockout show strongest asymmetric localization to one side of the cardiomyocyte nuclei (Figure 2A-E). This is followed by an intermediate asymmetry in *Lmna^+/-^*;*Lmnb1^-/-^*and a milder asymmetry in *Lmnb1^-/-^* cardiomyocytes (Figure 2A-E). This suggests that total lamin concentration, rather than lamin identity, sets the threshold for maintaining even distribution of these nuclear envelope structures^18,19^. Lamin-B1 appears to carry most of this function during embryogenesis when lamin-A expression is low^15,39^.

Unlike lamin-A knockouts^27,77^, *Lmnb1*^-/-^ cardiomyocytes show no increased DNA damage (Figure 3A, 3C-E), despite substantial disruption in nuclear and chromatin (judged by DAPI staining) organization (Figure 2F, 2G). This allowed us to analyze how lamin-B1 regulates genome interactions and gene expression during cardiomyocyte development without the confounding effects of DNA damage. The polarized NPC/LINC complexes together with chromosome elongation suggests a model in which cytoskeletal forces, transmitted through the NPC/LINC complexes, indirectly organize chromatin. Lamin-B1 may buffer these forces and keep NPCs/LINC complexes evenly distributed, and LADs at the nuclear periphery. As LAD undergo developmental remodeling in cardiomyocytes, lamin-B1 may be a node that modulates cytoskeletal forces and other regulatory processes to allow local formation of cardiomyocyte-specific LADs or non-LADs. When lamin-B1 is lost, the cytoskeletal forces pull on the remaining lamina asymmetrically, dragging NPCs, LINC complexes, and the chromosomes anchored at the periphery toward one side of the nuclei. The resulting chromosome territorial expansion would erode a subset of LADs as we observed for the partial or complete loss of some HiLands-B LAD regions. This would lead to rewired interactions between active (HiLands-R/O/Y) and repressive (HiLands-B/P) chromatin neighborhoods, providing a physical basis for the gene dysregulation we observe.

By P7, even though NPC, LINC, and microtubule organization is largely rescued in *Lmnb1^-/-^* cardiomyocytes, chromosome expansion and maturation defects persist (Figure 5F-H, S1, S3). This suggests that the embryonic window during which lamin-B1 modulates cytoskeletal force on chromatin leaves a durable imprint on 3D genome organization that cannot be unwound by increased lamin-A expression.

Cardiomyocytes undergo rhythmic contraction and a stereotyped maturation transition^36^, which can amplify the mechanical demand on the nuclear lamina^78^. The need for lamin-B1 to buffer cytoskeletal forces is therefore particularly visible in this cell type. However, the underlying principle by which lamin-B1 moderating mechanical input to preserve LADs and 3D genome organization may not be cardiomyocyte-specific. In cell types under lower mechanical strain, the same buffering function should still operate, but with subtler chromosome phenotypes that are difficult to discern at the whole chromosome level. Indeed, by studying cells such as mouse embryonic stem cells (mESC) that are under milder mechanical strain than cardiomyocytes, we showed that deleting all three lamins does not cause DNA damage, but results in a similar detachment of select HiLands-B LAD regions and altered chromatin neighborhoods, which can explain gene expression changes^55,79^.

Building on this physical role, our integrated Hi-C, HiLands, scRNA-seq, and motif analyses argue that lamin-B1 maintains LADs to gate the transcription-factor logic of maturation and cell-cycle exit in cardiomyocytes. Upon lamin-B1 loss, a fraction of the downregulated genes interacted more with repressive HiLands-P/B neighborhoods (Figure S6J, Table 4). This repressive environment may render binding motifs for Mef2d, Sp1, Nf1, E2F, and Banp less accessible and/or reduce the activities of these transcription factors, thereby compromising cardiomyocyte structural maturation and cell-cycle exit (Figure 7A, 7B, Table 5)^67,68,70–72^. Conversely, a fraction of the upregulated genes lose contact with repressive HiLands-B and can become more permissive for Sp1, Banp, Ronin, HiNF-P, and Nrf1 to bind and transcribe their target genes (Figure 7A, 7B, Table 4, Table 5)^68,69,73–75^. We showed the example of how the dysregulation of Cdk4, Jun, and E2F3 can propagate these chromatin-neighborhood effects through downstream transcriptional cascades (Figure 7C-F, Table 4) ^49,70,76^. By focusing on lamin-B1 in cardiomyocytes, we therefore establish a LAD-gated logic that links a ubiquitously expressed nuclear lamina protein to the transcription factors known to drive cardiomyocyte maturation and halt division. Additionally, if lamin-B1-dependent LADs help enforce cardiomyocyte cell-cycle exit, then transient and controlled perturbation of this lamin could become a candidate strategy to reopen the cell cycle in adult cardiomyocytes for regenerative purposes.

More broadly, lamin-B1 levels have been found to decline in different cell types upon aging^80–84^ and in oncogene induced senescence^85^. Our findings suggest that LAD remodeling under reduced lamin-B1 may not merely be a passive consequence of these states but an active enabler that shifts proliferation- and survival-gene promoters out of repressive lamina-tethered neighborhoods and into active interior compartments, where transcription factors can drive senescence escape. Conversely, the same logic implies that stabilizing lamin-B1 or its LAD anchoring could resist these transitions.

### Limitations of the Study

Several technical limitations constrain the functional experiments that could further test these findings. While we are able to culture primary embryonic and early neonate mouse cardiomyocytes, extensive culture results in progressive dedifferentiation. Furthermore, primary mouse cardiomyocytes have well-documented low transfection efficiency through traditional means. These collectively limit our ability to perform rescue experiments where we reintroduce or overexpress lamins, as well as genomic manipulation for chromatin tethering experiments or deletions. Additionally, the slow turnover of lamin-B1 presents a barrier to test acute loss of lamin-B1 in postnatal cardiomyocytes. Future studies using viral delivery systems and auxin-inducible degron approaches to acutely deplete lamin-B1 will be important to further examine our findings.

## Author Contributions

K.A.B. performed experiments, bioinformatic and image analysis, and visualization. X.Z. advised on bioinformatic analyses. Y.K., L.K., and R.M. developed low-cell Hi-C methodology. Y.Z. and K.A.B. conceptualized the study, wrote and edited the manuscript. Y.Z. and Y.K. supervised aspects of the work and acquired funding.

## Acknowledgements

We thank Allison Pinder, Joseph Tran, Fred Tan, and Javier Carpinteyro Ponce for assistance with sequencing; Mahmud Siddiqi for microscopy and image analysis; Lynne Hugendubler for technical support; and Joseph Tran, Sara Debic, Wesley Yon, and Ross Pederson for valuable technical advice and feedback. We also thank members of the Zheng lab for helpful discussions. This work was supported by grants R01GM106023 (Y. Zheng, R.D. Goldman), R01GM110151 (Y. Zheng), 1R01GM157598-01 (Y. Zheng), RS-2023-00222784 (Y. Kim), RS-2025-25441283 (Y. Kim), and RS-2024-00437643 (Y. Kim).

## Declaration of Interests

The authors declare no competing interests.

## Declaration of AI

During preparation of the manuscript, Y.Z. and K.A.B. utilized ChatGPT-5 mini (OpenAI) and Opus 4.7 (Claude) to assist with editing sections of the text for clarity, which was subsequently reviewed and edited by the authors. Y.Z. and K.A.B. utilized Zo Computer (https://www.zo.computer) to aid in literature review.

## Methods

## RESOURCE AVAILABILITY

### Lead Contact

Requests for additional information should be sent to the corresponding authors, Katherine Bossone and Yixian Zheng.

### Materials availability

Lamin and Cre mouse lines used to generate the genotype combinations in this study are available from Jax and Charles River (see Methods for strain details).

### Data and code availability

All bulk RNA-seq, scRNA-seq, CUT&RUN, ATAC-seq, and Hi-C datasets generated in this study have been deposited in GEO as a SuperSeries with the accession number GSE330298, which contains the accession numbers GSE330013, GSE330099, GSE330100, GSE330102, and GSE330103. Custom code created and used in this study will be publicly available at https://github.com/katherinebossone/Lamin_CM_Paper.

## KEY RESOURCES TABLE

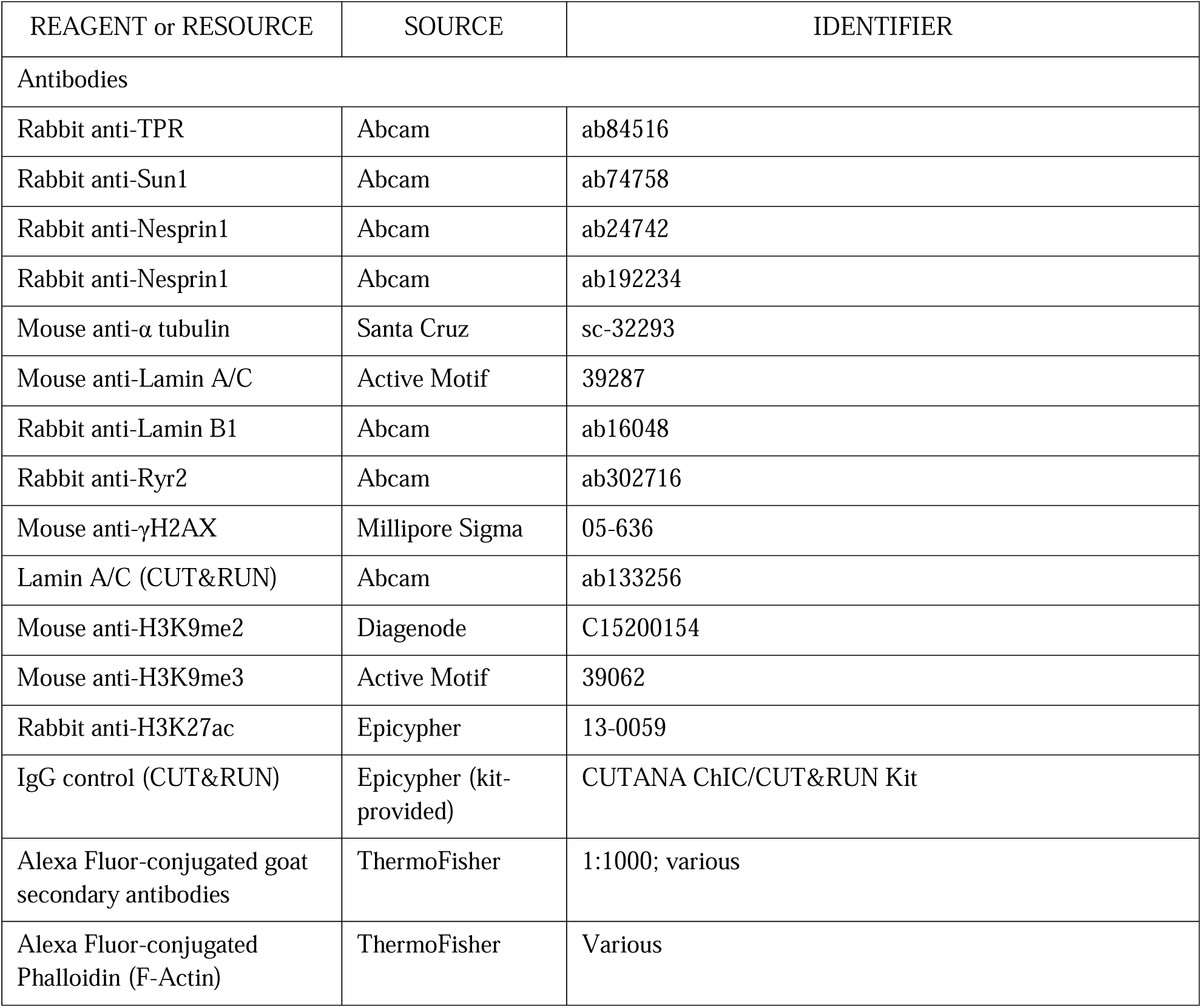
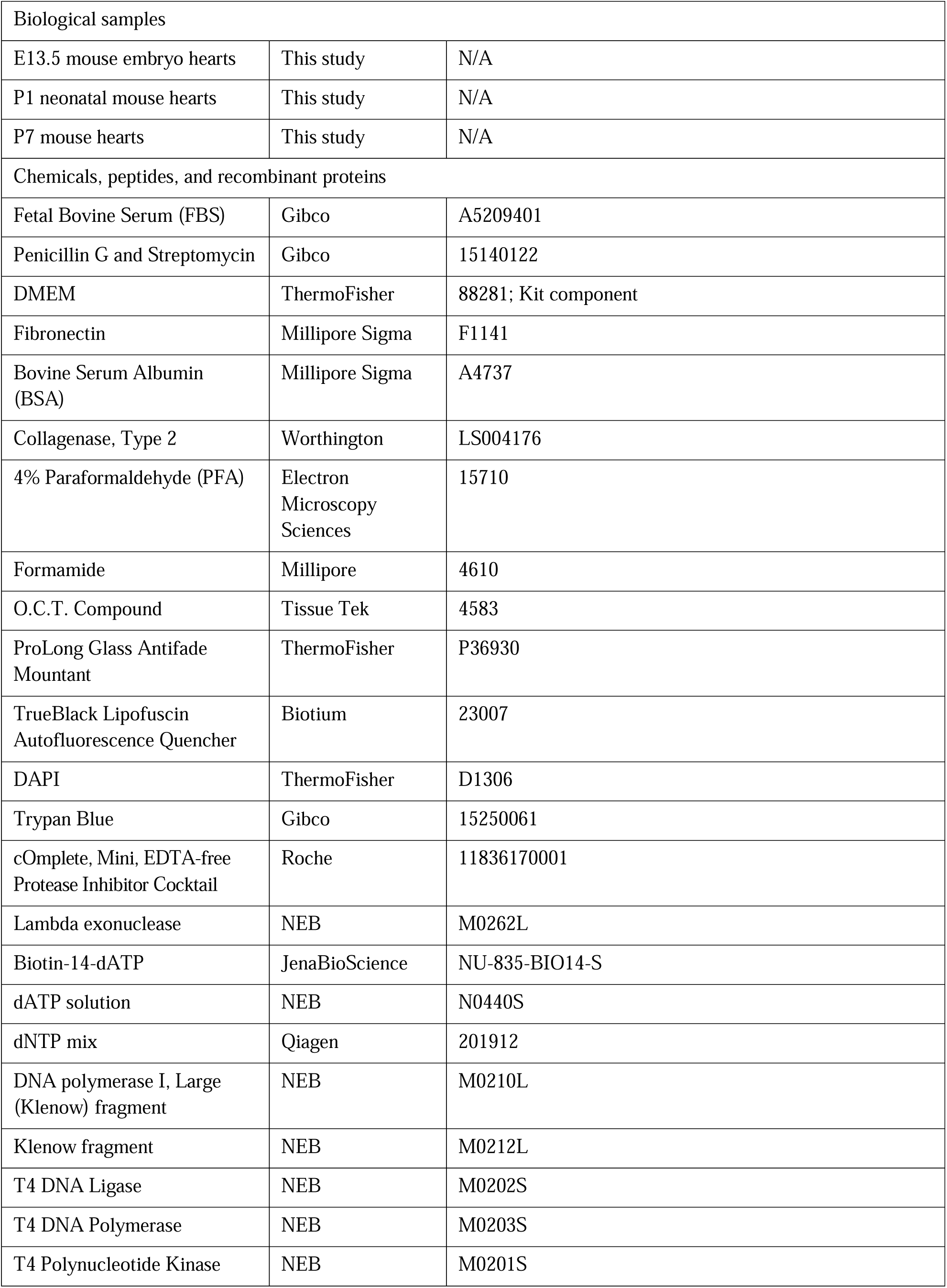
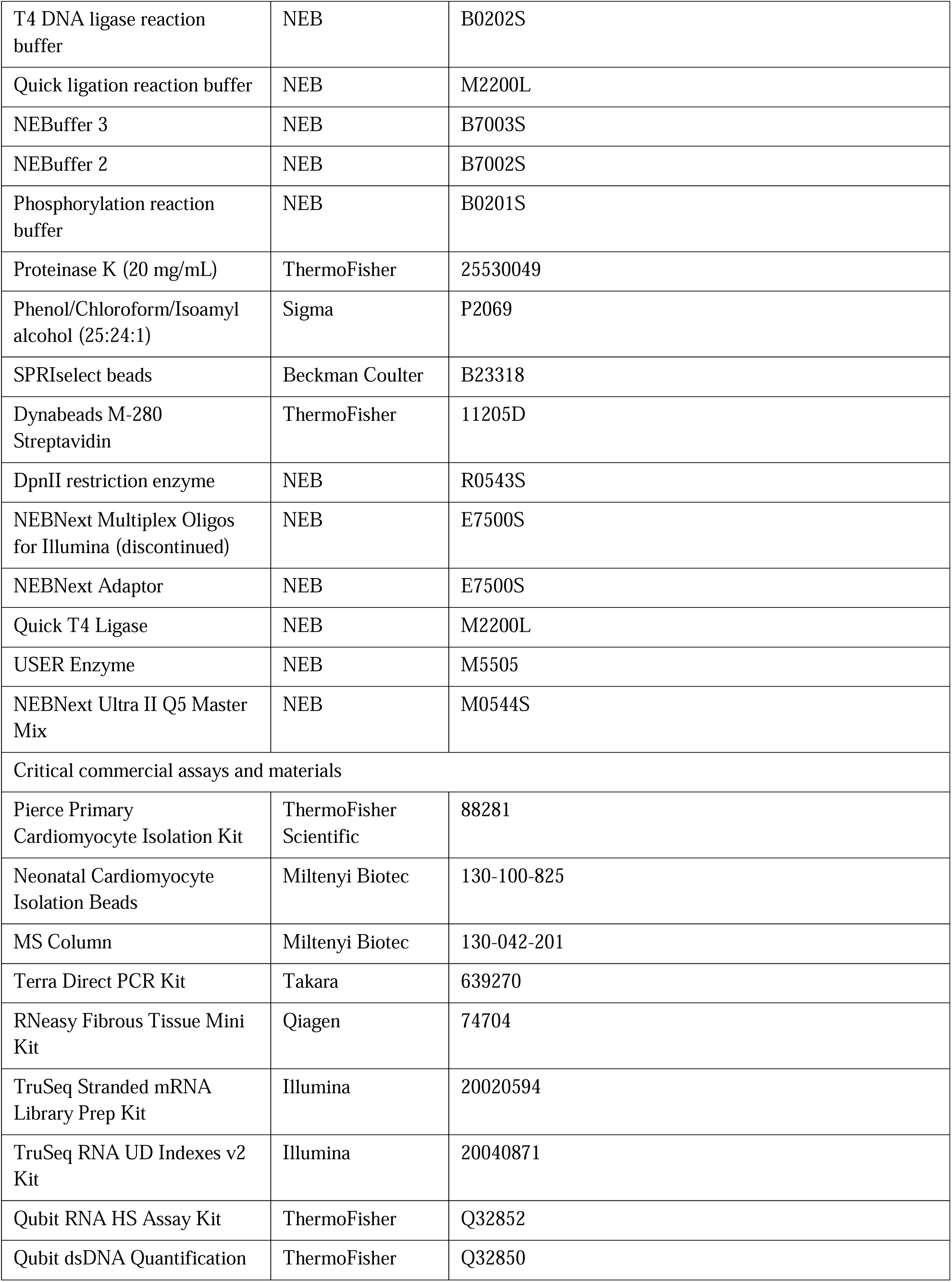
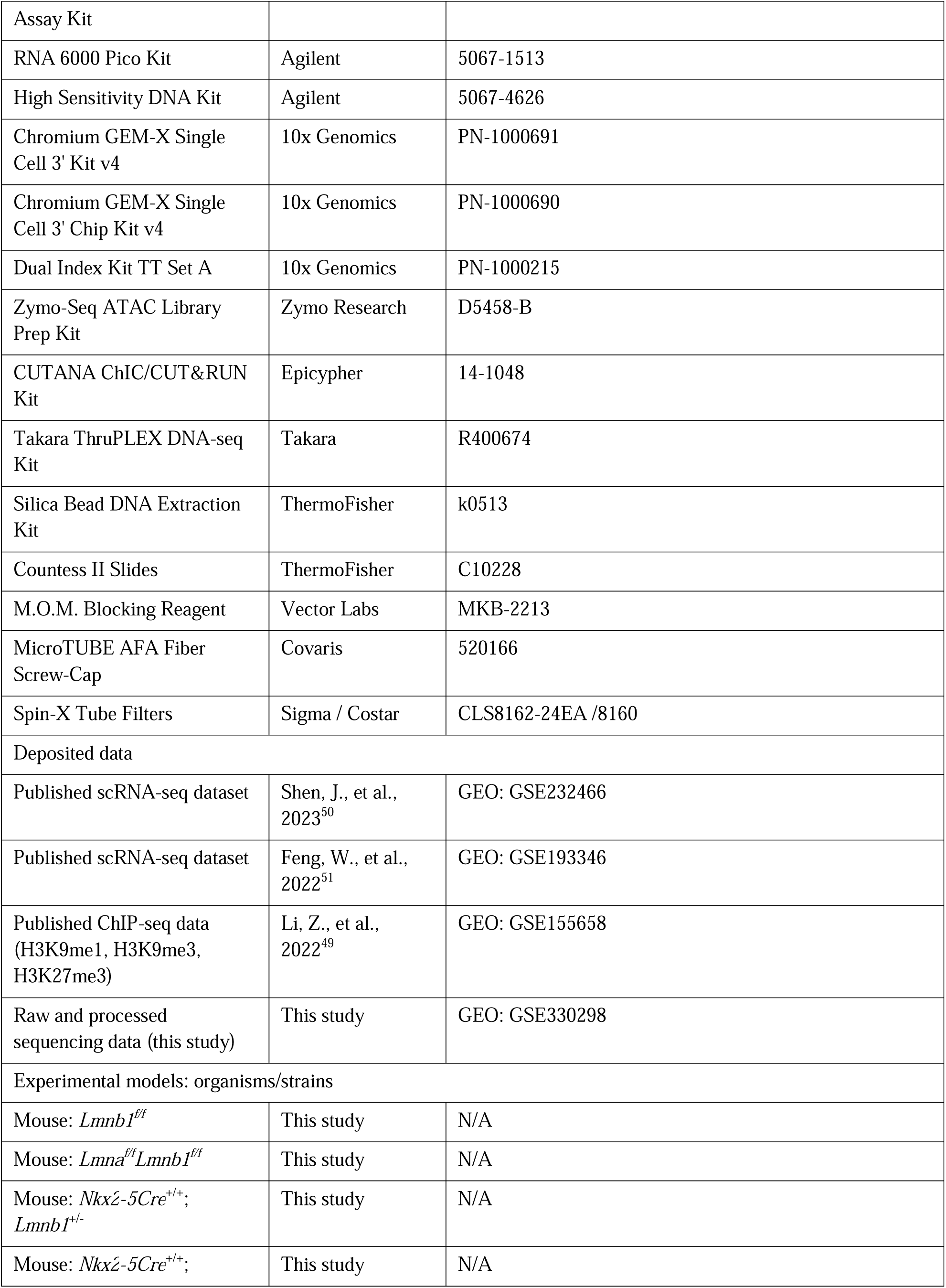
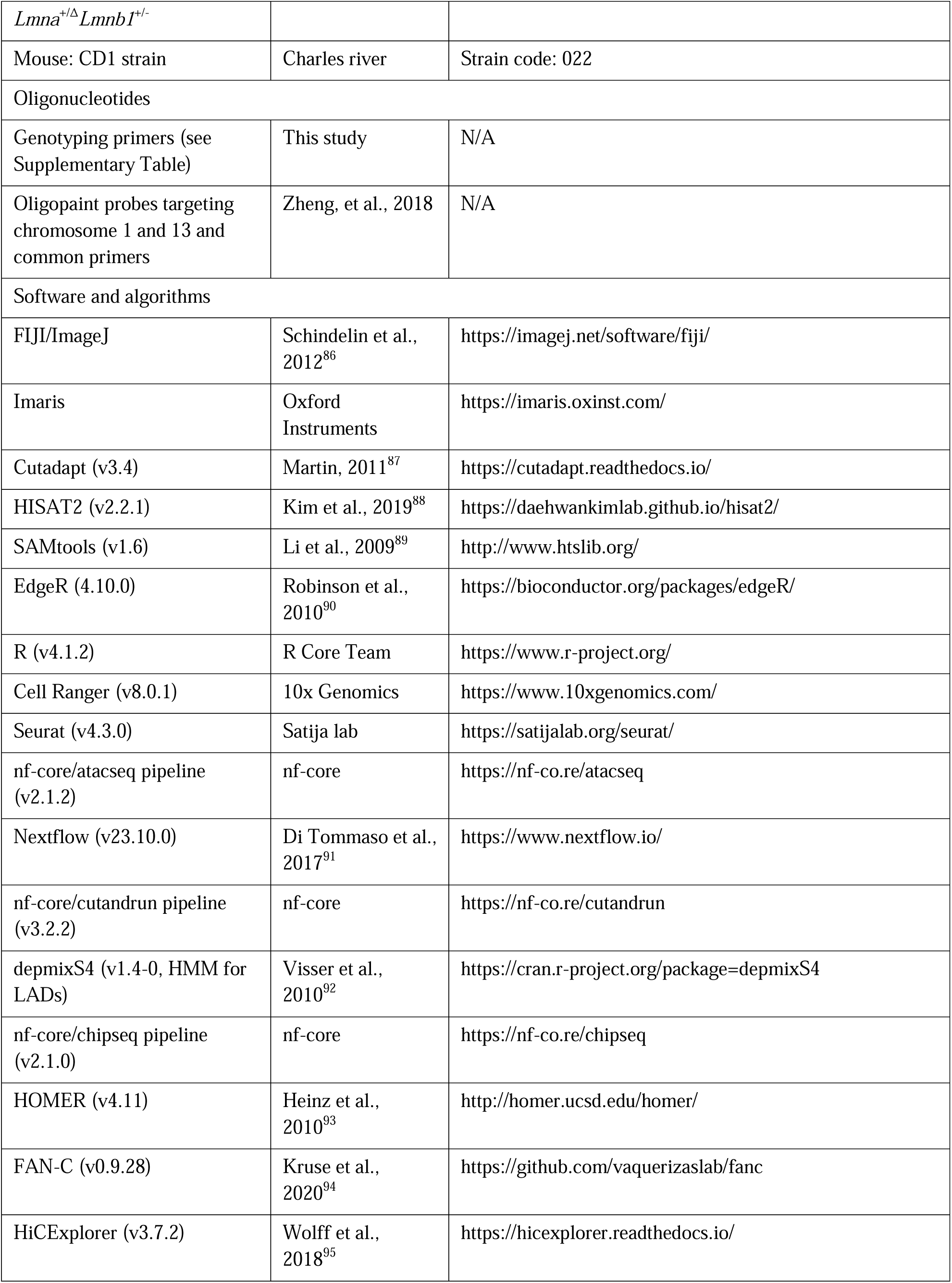
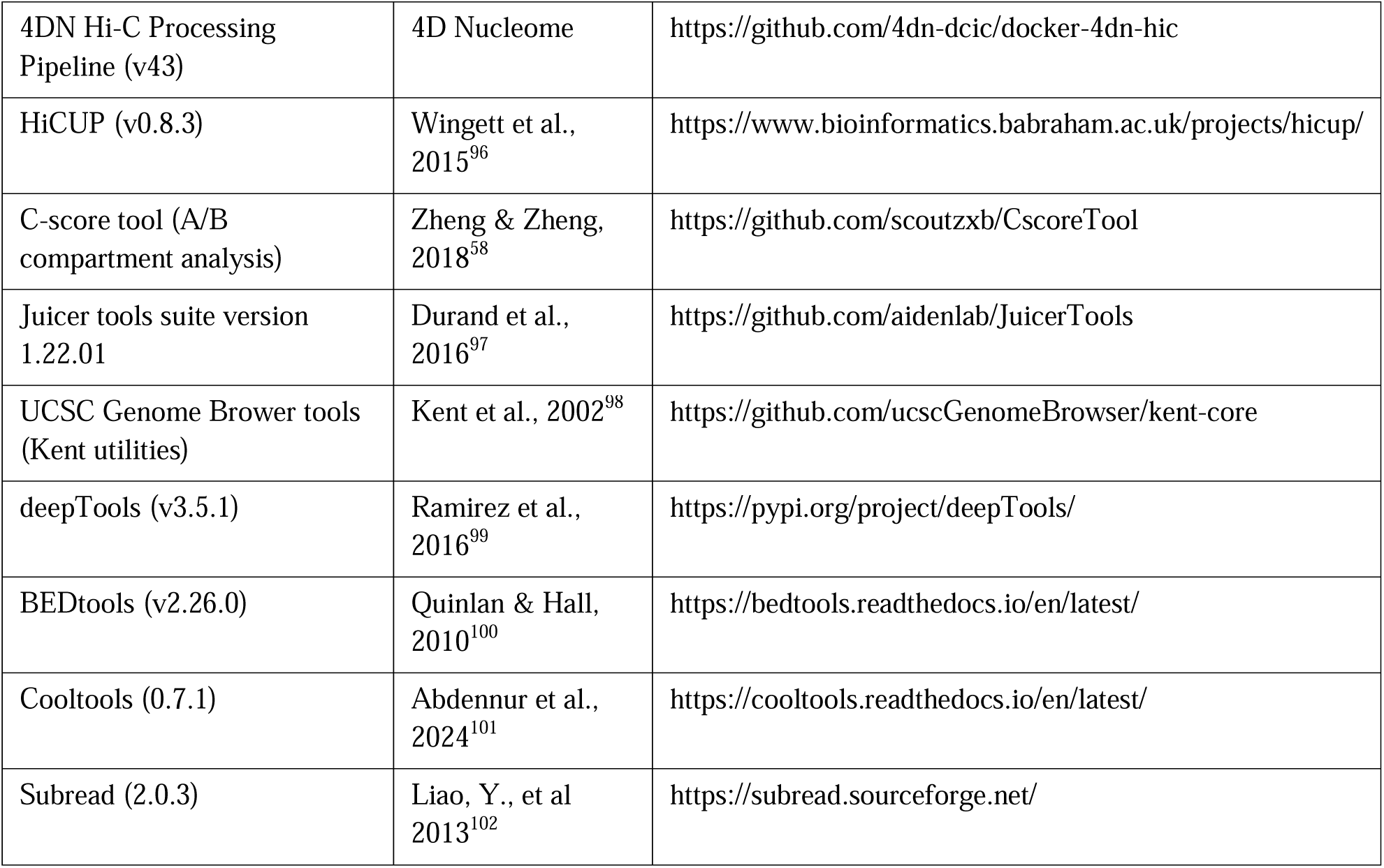

## EXPERIMENTAL MODEL AND SUBJECT DETAILS

### Animals

All animal procedures were approved by and performed in accordance with the guidelines of Carnegie’s Institutional Animal Care and Use Committee (IACUC) and the NIH. Mice were housed in a semi-permeable mouse facility with 12-hour light/dark cycles, a controlled temperature of 20-22°C, and humidity of 30-50%. Cages were regularly cleaned, and clean food and water were provided. Animal health was monitored using a sentinel surveillance system, in which a designated sentinel cage is exposed to soiled bedding from colonies within the rack. Sentinel mice are regularly tested for pathogens at Charles River Laboratories. Colony mice are further monitored for physical injuries and visible illness.

### Mouse strains

To produce the various lamin-A and -B1 knockout combinations, conditional alleles for *Lmna* and *Lmnb1* previously generated in our lab were utilized. *Lmnb1^f/f^* mice were produced with the *Lmnb1*^tm1a(EUCOMM)Wtsi^ allele (EUCOMM project, International Mouse Strain Resource)^13^. A *Lmna^f/f^*;*Lmnb1^f/f^*line was generated by crossing *Lmnb1^f/f^* mice with *Lmna^f/f^* mice (JAX stock #026284), which carriess *loxP* sites flanking exon 2 of *Lmna*^103^. *Nkx2-5Cre^+/+^*;*Lmna^+/-^*;*Lmnb1^+/-^*and *Nkx2-5Cre^+/+^*;*Lmnb1^+^*^/-^ lines were generated by crossing mice heterozygous for the *Lmna* and *Lmnb1* alleles generated previously with mice homozygous for Cre recombinase driven by the Nkx2-5 promoter (JAX stock #030047). Mice carrying the Nkx2-5Cre transgene and heterozygous for lamin-A and lamin-B1, or lamin-B1 alone (*Nkx2-5Cre^+/+^*;*Lmna^+/-^*;*Lmnb1^+/-^*and *Nkx2-5Cre^+/+^*;*Lmnb1^+^*^/-^), were crossed with mice homozygous for the floxed lamin-A and/or lamin-B1 alleles (*Lmna^f^*^/f^;*Lmnb1^f/f^*and *Lmnb1^f/f^*). These breedings produced the following genotypes: *Nkx2-5Cre^+/-^*;*Lmna^+^*^/-^;*Lmnb1^+^*^/-^, *Nkx2-5Cre^+/-^*;*Lmna*^-/-^;*Lmnb1^+^*^/-^, *Nkx2-5Cre^+/-^*;*Lmna^+/-^*;*Lmnb1^-^*^/-^, *Nkx2-5Cre^+/-^*;*Lmna*^-/-^;*Lmnb1^-^*^/-^, *Nkx2-5Cre^+/-^*;*Lmnb1^+^*^/-^, and *Nkx2-5Cre^+/-^*;*Lmnb1^-^*^/-^ (Figure 1A). All conditional knockout and Cre-driver lines were maintained on a mixed CD1, 129Sv, and C57BL/6J genetic background. Sex was not treated as a biological variable, and both sexes were included in image analyses of cytoskeletal and nuclear organization. Sequencing data were derived solely from male mice to remove sex-dependent effects, and littermates were prioritized to minimize batch effects. For lamin allele knockout experiments, either *Lmnb1^+^*^/-^ or *Lmna^+/^*^Δ^;*Lmnb1^+^*^/-^ cardiomyocytes were used as controls, while wild-type cardiomyocytes for all other experiments were collected from CD1 mice.

## METHOD DETAILS

### Cardiomyocyte isolation and culture

For E13.5 cardiomyocytes, pregnant mice were euthanized by cervical dislocation, and embryos were dissected and kept on ice during genotyping. Hearts were dissected, minced, and dissociated using the Pierce Primary Cardiomyocyte Isolation Kit (ThermoFisher Scientific, 88281). Briefly, the cardiomyocyte isolation enzyme was prepared according to the manufacturer’s instructions, and hearts were dissociated at 37°C for 15 minutes. Hearts were washed in Hank’s Buffered Saline Solution (HBSS) and dispersed into single cells by pipetting in cardiomyocyte culture medium consisting of 10% (volume/volume) Fetal Bovine Serum (FBS; Gibco) and 1% (volume/volume) Penicillin G/Streptomycin (Pen/Strep) in DMEM. For P1 cardiomyocytes, neonates were decapitated and hearts were dissected and processed as for E13.5 samples, except that enzymatic dissociation was extended to 25 minutes.

For culturing dissociated E13.5 cardiomyocytes, glass coverslips were sterilized in isopropanol, placed in a 24-well plate, and washed with HBSS. Coverslips were then coated with 5 μg/mL fibronectin (Millipore Sigma, F1141) in HBSS for 2 hours at 37°C. After coating, coverslips were washed with HBSS and kept in cardiomyocyte culture medium until cell seeding. Cells were seeded at approximately 100,000 cells/well in a 24-well plate and incubated overnight (37°C, 5% CO2, 90-95% humidity). The following day, the culture medium was refreshed.

For purification of P1 cardiomyocytes, dissociated cells from the hearts were pelleted at 300 x g for 5 minutes and resuspended in MACS buffer consisting of 0.5% (weight/volume) bovine serum albumin (BSA) and 2mM EDTA in phosphate-buffered saline (PBS). The cells were then incubated with neonatal cardiomyocyte isolation magnetic beads (Miltenyi Biotec, 130-100-825) for 15 minutes while rotating at 4°C in the dark; these beads bind non-cardiomyocyte cells. Samples were brought up to 500 μL with MACS buffer and run through an MS column (Miltenyi Biotec, 130-042-201). The flowthrough containing enriched cardiomyocytes was collected. MS columns were rinsed twice with 500 μL of MACS buffer to recover additional cardiomyocytes. The combined cardiomyocytes were pelleted at 300 x g for 5 minutes and resuspended in PBS. Viability and cell concentration were assessed using the Countess II Automated Cell Counter (Fisher Scientific) with the associated counting slides (ThermoFisher Scientific, C10228). The cardiomyocyte suspension was diluted 1:1 with Trypan Blue to detect and count dead cells. This achieved a cardiomyocyte purity of >80%.

### Genotyping

Mouse embryos were genotyped using both limb and tail tissues, while neonatal and adult mouse samples were genotyped from tail biopsies only. Genotyping samples were digested in 500 μL of 50mM sodium hydroxide and incubated at 90°C for 7 minutes. After digestion, sodium hydroxide was neutralized with 50 μL of 10 mM Tris-HCl (pH 8.0), and samples were centrifuged at 20,817 x g for 4 minutes at room temperature. 1 μL of supernatant was used as template DNA for PCR with the Terra PCR kit according to the manufacturer’s instructions. Cycling conditions were 98°C for 2 minutes, followed by 35 cycles of 98°C for 10 seconds, 63°C for 15 seconds, and 68°C for 1 minute. PCR products were visualized on 2% agarose gels. Genotyping primers and expected band sizes are listed in supplementary table 6.

### Histology

For E13.5 samples, whole embryos were dissected and fixed in 4% paraformaldehyde (PFA) in PBS at 4°C for 24 hours. For P1 and P7 time points, hearts were dissected prior to fixation in 4% PFA in PBS and stored under the same conditions for 24 hours. P7 tissues designated for fluorescence in situ hybridization (FISH) were instead fixed for 4 hours, as over-fixation impaired labeling efficiency. After fixation, all tissues were transferred to 30% sucrose in PBS and incubated at 4°C for 24 hours. Samples were then embedded in O.C.T. compound (Tissue Tek) and stored at -80°C until sectioning.

### Immunofluorescence

For whole-heart and embryonic tissues, samples were equilibrated to -23°C and sectioned at 12-20 μm thickness on a Leica CM3050 S cryostat. Sections were collected onto Superfrost Plus microscope slides and air-dried overnight at room temperature. Prior to staining, sections were rehydrated in PBS for 5 minutes. Sections were permeabilized for 25 minutes in 0.25% (volume/volume) Triton X-100 in PBS and briefly rinsed with PBS. Sections were then blocked for 1 hour in blocking solution containing 10% (weight/volume) bovine serum albumin (BSA) and 10% (volume/volume) normal goat serum in PBS. For sections stained with mouse primary antibodies, M.O.M. blocking reagent (Vector Labs, MKB-2213) was used according to the manufacturer’s instructions prior to antibody incubation.

For P7 Ryr2 staining, tissue sections were partially dissociated prior to rehydration to reduce cell overlap and facilitate visualization of Ryr2 distribution in individual cardiomyocytes. Sections were incubated with 1 mg/mL Collagenase type II (Worthington, LS004176) in PBS for 30 minutes at room temperature, then mechanically dispersed into a near-single-cell solution by gentle pipetting. The cell suspension was stained as above in solution, spinning down at 300g for 5 minutes after each solution swap. After staining they were resuspended in mounting media and dispensed onto slides with coverslips.

For isolated cardiomyocyte cultures, cells were fixed for 10 minutes in 4% PFA in PBS, washed three times with PBS, and permeabilized for 10 minutes in 0.25% Triton X-100 in PBS. Cells were then washed with PBS and blocked for 1 hour in blocking solution containing 10% BSA and 10% normal goat serum in PBS. For all microtubule immunostaining, PBS was substituted with BRB80 buffer (80mM PIPES,1mM MgCl2, 1mM EGTA, pH 6.8) throughout the protocol to preserve microtubule structure.

For immunostaining of both tissue sections and cultured cardiomyocytes, samples were incubated with primary antibodies diluted in blocking solution either overnight at 4°C or for 2 hours at room temperature (see Antibodies section). Samples were then washed three times with PBS for 5 minutes each and incubated with the appropriate secondary antibodies and DAPI (5μg/mL; ThermoFisher, D1306) diluted in blocking solution for 2 hours at room temperature, protected from light. Samples were washed three additional times in PBS and mounted with ProLong Glass Antifade Mountant (ThermoFisher, P36930). To reduce background fluorescence in tissue sections, sections were treated with TrueBlack Lipofuscin Autofluorescence Quencher (Biotium, 23007) for 30 seconds, followed by a PBS rinse, prior to mounting. Images were acquired on a Leica SP5 confocal microscope.

### Antibodies

The primary antibodies used in this study were as follows: rabbit anti-TPR (Abcam, ab84516; 1:200), rabbit anti-Sun1 (Abcam, ab74758; 1:200), rabbit anti-Nesprin1 (Abcam, ab24742; 1:200 and Abcam, ab192234; 1:200), mouse anti-α-tubulin (Santa Cruz, sc-32293; 1:1,000), mouse anti-lamin-A/C (Active Motif, 39287; 1:400), rabbit anti-lamin-B1 (Abcam, ab16048; 1:400), rabbit anti-Ryr2 (Abcam, ab302716; 1:200), and mouse anti-γH2AX (Millipore Sigma, 05-636; 1:1,000). Alexa Fluor-conjugated goat secondary antibodies (ThermoFisher) were used at 1:1,000. To label F-actin, Alexa Fluor-conjugated phalloidin (ThermoFisher) was used at 33nM.

### Quantification of nuclear polarization

Cardiomyocytes were identified by phalloidin staining of striated F-actin. For each cardiomyocyte nucleus, a line was drawn across the long axis of the nucleus starting at the point of highest fluorescence intensity. To account for variation in nuclear length, the line was divided into 10 equal segments, and the average fluorescence intensity within each segment was calculated. Each profile was then normalized to the section of maximum intensity, allowing comparison of polarization between genotypes and replicates done on different days. Figure 2C illustrates how this quantification was performed.

## Quantification of nuclear morphology and DNA damage

For each E13.5 image, thresholding was applied to DAPI-stained nuclei until the nuclear areas were fully included. Nuclei that failed to be clearly defined, or that did not exhibit a striated actin appearance in the surrounding cytoplasm, were excluded from analysis. The nuclei were then measured using the Shape Descriptors function in Fiji (ImageJ), defined as (4 × area)/(π × major axis^2^) to assess their roundness. For DNA damage analysis, the nuclear area defined by DAPI were applied to the corresponding γH2AX channel, and the mean γH2AX fluorescence intensity per nucleus was calculated.

For each P1 image, due to the difficulty of thresholding over background in tissue sections, percent of nuclei that were γH2AX positive was calculated. γH2AX-positive nuclei were defined by nuclei with greater than ≥2 foci.

### Fluorescence In Situ Hybridization (FISH)

#### 1. Probe preparation

Oligopaint probes targeting the entirety of chromosomes 1 and 13 were prepared from libraries designed and described in our previous study^55^. Briefly, 1 ng of each chromosome library was amplified with the corresponding common primers conjugated to Alexa Fluor 488. DNA was precipitated by adding to the PCR product an ice-cold solution of 4 M ammonium acetate (1/10 volume), 100% ethanol (2.25 volumes), and 20 mg/mL glycogen (1/50 volume), with all volumes calculated based on the PCR product volume. Pellets were washed with 2.25 volumes of ice-cold 70% ethanol and air-dried at 42°C until visibly dry. Probes were resuspended in 50 μL of deionized water and treated with 1 μL of lambda exonuclease (NEB, M0262L) to digest residual single-stranded DNA. DNA was precipitated again as described above and resuspended in deionized water. Final probes were quantified using a NanoDrop spectrophotometer (yields of 30-50 pmol/μL) and verified by gel electrophoresis, with a single band visible at approximately 88 bp.

#### 2. FISH on isolated cardiomyocyte culture

Cardiomyocytes on coverslips were fixed for 10 minutes in 4% PFA in PBS at room temperature and permeabilized in 0.25% Triton X-100 in PBS at room temperature for 10 minutes. Cells were then incubated in 20% (volume/volume) glycerol in PBS overnight at room temperature. The following day, coverslips were dipped in liquid nitrogen (∼5 seconds) and thawed on a paper towel; this freeze-thaw cycle was repeated five times to further permeabilize the cells. Coverslips were washed once in PBS for 1 minute, followed by two additional 5-minute PBS washes. Cells were then treated with 0.1 M HCl in water for 5 minutes to denature proteins and depurinate DNA, improving probe access. Coverslips were washed three times for 1 minute each in 2X Saline-Sodium Citrate (SSC) buffer at room temperature. Coverslips were pre-equilibrated in 2X SSC containing 50% (volume/volume) formamide for 4 hours at 37°C. During this time, 40 pmol of probe per 25 μL reaction was added to hybridization buffer (2X SSC, 10% (weight/volume) dextran sulfate, 50% (volume/volume) formamide) and pre-warmed at 37°C for 2 hours. The probe-containing hybridization solution was pipetted onto Superfrost Plus microscope slides, and coverslips were placed cell-side down onto the solution, sealed with rubber cement, and denatured at 80°C on a heat block for 3 minutes. Slides were then transferred to a humidified box and incubated at 37°C for 14 hours. After hybridization, coverslips were removed and cells were washed twice in 2X SSCT (2X SSC, 0.1% (volume/volume) Tween-20) at 60°C for 15 minutes per wash. This was followed by one 10-minute wash in 2X SSCT and one 10-minute wash in 0.2X SSCT, both at room temperature. Coverslips were then briefly washed in 1X SSCT, after which immunofluorescence staining for TPR was performed as described above, with 1X SSCT replacing PBS in all wash and incubation steps.

#### 3. FISH on tissue sections

FISH on tissue sections was performed as previously described^104^. Briefly, P7 hearts in O.C.T. compound were equilibrated to -23°C, sectioned at 20 μm, and air-dried at room temperature overnight. Tissue sections were rehydrated in sodium citrate buffer (10 mM sodium citrate, 0.05% (volume/volume) Tween-20, pH 6.0) for 5 minutes at room temperature. To improve probe penetration, tissue sections were further incubated in sodium citrate buffer for 25 minutes at 80°C, followed by 10 minutes of cooling at room temperature. Slides were removed from the sodium citrate buffer, and a hydrophobic barrier was drawn around each tissue section using a PAP pen. Sections were washed twice with 2X SSC for 5 minutes each at room temperature; during this wash, pedestals (#1.5 coverslips adhered with nail polish and allowed to dry) were arranged adjacent to each section. Slides were then transferred into 50% (volume/volume) formamide in 2X SSC and incubated for 4 hours at 37°C. During this time, 40 pmol of probe per 25 μL reaction was added to hybridization buffer (composition as above) and pre-warmed at 37°C for 2 hours. Slides were drained, the probe solution was applied to each section, and a coverslip was placed on top of the pedestals and sealed with rubber cement. Sections were denatured at 80°C for 5 minutes, transferred to a humidified chamber at 37°C, and allowed to hybridize for 3 days. After hybridization, coverslips were removed and slides were washed three times in 2X SSC for 15 minutes each at room temperature, followed by two washes in 0.1X SSC at 60°C for 5 minutes each, and a 2-minute equilibration in 2X SSC at room temperature.

#### 4. Imaging and volume quantification

Confocal images were acquired on a Leica SP5 confocal microscope equipped with a 63X/1.4 NA oil objective. For each set of experiments, image stacks were acquired with a z-step of 126 nm using identical laser settings. Image files were imported into Imaris (Bitplane) to generate 3D reconstructions. Random images from lamin-B1 control and knockout samples were used to train the Imaris machine-learning segmentation tool until chromosomes were well resolved across most images. Minor manual corrections were performed in regions where chromosome identification was inaccurate. Nuclei with poorly defined chromosomes, such as those with high background signal, were excluded from downstream analyses.

### Bulk RNA-sequencing (RNA-seq)

Bulk RNA-seq was performed on whole P7 hearts using the RNeasy Fibrous Tissue Mini Kit (Qiagen, 74704) according to the manufacturer’s instructions. Whole P7 hearts were dipped in liquid nitrogen for 10 seconds and crushed with a mortar and pestle. Crushed tissue was placed in 300 μL of Buffer RLT supplied by the kit containing 10 μL of β-mercaptoethanol, 590 μL of RNase-free water and 10 μL of supplied proteinase K were added to each sample, which was then incubated at 55°C for 10 minutes. Samples were centrifuged at 10,000 x g for 3 minutes, and the supernatant was collected. 450 μL of 100% ethanol was added to each sample and mixed by pipetting. Samples were applied to the provided RNA collection columns and washed with Buffer RW1. 10 μL of supplied DNase was added to each column and incubated at room temperature for 15 minutes. Columns were then washed once with Buffer RW1, followed by two washes with Buffer RPE. RNA was eluted from each column with 20 μL of RNase-free water; the eluate was passed through the column once more to recover additional RNA. RNA was quantified using the Qubit RNA HS Assay Kit (ThermoFisher, Q32852), and quality was verified using the 2100 Bioanalyzer System with the RNA 6000 Pico Kit (Agilent, 5067-1513). Poly-A-selected libraries were prepared with the TruSeq Stranded mRNA Library Prep kit (Illumina, 20020594) and TruSeq RNA UD Indexes v2 (Illumina, 20040871). Final libraries were quantified with the Qubit dsDNA Quantification Assay Kit (ThermoFisher, Q32850), and library quality was verified on the 2100 Bioanalyzer System using the High Sensitivity DNA Kit (Agilent, 5067-4626). Sequencing was performed on the Illumina NextSeq 500 with 75 x 75 bp single-end reads.

### Analysis of bulk RNA-seq

Reads were adapter- and quality-trimmed using Cutadapt, removing 10 bases from the 5′ end and 5 bases from the 3′ end. Trimmed reads were aligned to the mouse mm10 reference genome using HISAT2. The resulting alignments were converted to BAM format, sorted, and indexed using SAMtools. The Subread (v2.0.3) package FeatureCounts was used to generate gene-level read counts from aligned reads using the mm10 annotation. Differential expression analysis was performed in R using the edgeR package. Genes with fewer than 10 reads in at least 4 samples were excluded from analysis. A cutoff of |Log_2_FC| > 0.26 and p < 0.05 was used to identify differentially expressed genes.

### Single-cell RNA-sequencing (scRNA-seq) of neonatal hearts

P1 hearts from one *Lmnb1^+/-^* and two *Lmnb1^-/-^* littermate pups were dissociated separately as described above. Cells were pelleted at 300 x g for 5 minutes and resuspended in PBS. 10X libraries were prepared with the Chromium GEM-X Single Cell 3’ Kit v4 (10x Genomics, PN-1000691), Chromium GEM-X Single Cell 3’ Chip Kit v4 (10x Genomics, PN-1000690), and Dual Index Kit TT Set A (10x Genomics, PN-1000215) according to the manufacturer’s protocol (CG000731). dsDNA was quantified with the Qubit dsDNA Quantification Assay Kit, and library quality was verified on the 2100 Bioanalyzer System with the High Sensitivity DNA Kit. Sequencing was performed on the Illumina NextSeq 1000 according to the manufacturer’s recommendations.

### Analysis of scRNA-seq

Reads were processed using Cell Ranger (v8.0.1) following the 10x Genomics protocol. The Mus musculus reference genome (mm10, version 3.0.0) was used for alignment and gene quantification. Downstream analyses were performed in R (v4.1.2) using the Seurat package (v4.3.0) following standard workflows from the Satija lab. Gene-cell count matrices from three samples were imported and converted into Seurat objects. Cells expressing fewer than 100 genes or more than 8,000 genes, as well as cells with greater than 30% mitochondrial transcript content, were excluded from further analysis. Filtered datasets were merged and log-normalized, and highly variable features were identified, followed by scaling and principal component analysis (PCA). To improve normalization and integration across samples, SCTransform was applied, and datasets were integrated using canonical correlation analysis (CCA)-based integration. Dimensionality reduction was performed using PCA, and the first 17 principal components were used to construct a shared nearest-neighbor graph. Cells were clustered at a resolution of 0.5 and visualized using Uniform Manifold Approximation and Projection (UMAP). Rdata files from previously published datasets were downloaded and preprocessed using the same workflow prior to integration.

Cell-type populations were annotated based on the expression of canonical marker genes, as previously described^51,54^. Datasets were integrated both with all cell populations included, to assess overall cardiac population composition, and with non-ventricular cardiomyocyte populations excluded, to enable focused analyses of cardiomyocyte-specific transcriptional programs. For the cardiomyocyte-focused analysis, each filtered dataset was independently normalized using SCTransform to account for technical variation. Integration anchors were identified across the two previously published datasets and our datasets using Canonical Correlation Analysis (CCA) with the first 12 principal components, and these anchors were used to integrate the datasets into a single combined object. The integrated dataset was then scaled, and PCA was performed using the same 12 dimensions. A shared nearest-neighbor graph was constructed using these principal components, and unsupervised clustering was performed at a resolution of 0.5. For visualization, UMAP was applied using the first 12 principal components. All downstream analyses were conducted on the SCTransform (SCT) assay of the integrated object. Compact cardiomyocyte clusters (clusters 0-4 and 6-8) were combined into one cluster and differential gene expression analysis was performed using FindMarkers. The gene analysis was limited to genes expressed in more than 25% of cells in at least one genotype, with significance thresholds set at |Log_2_FC| > 0.26 and p < 0.05.

### Assay for Transposase-Accessible Chromatin sequencing (ATAC-seq) and processing

P1 cardiomyocytes were isolated and purified as described above, and an aliquot of 50,000 cells per sample was used for ATAC-seq. ATAC-seq libraries were prepared using the Zymo-Seq ATAC Library Prep Kit (Zymo, D5458-B) according to the manufacturer’s protocol. Briefly, live cells were lysed and nuclei were isolated using the kit-supplied reagents. Nuclei were incubated with the provided transposition mix at 37°C for 30 minutes, and DNA was purified using the supplied Zymo-Spin IC Columns. Sequencing libraries were prepared using the kit-provided library reagents and amplified with 10 cycles of PCR. Libraries were quantified with the Qubit dsDNA Quantification Assay Kit, and library quality was verified on the 2100 Bioanalyzer System with the High Sensitivity DNA Kit. Sequencing was performed on the Illumina NextSeq 1000 with 50/50 bp paired-end reads.

### ATAC-seq analysis

Samples were processed using the nf-core/atacseq pipeline (v2.1.2) with Nextflow (v23.10.0). The pipeline performed adapter trimming, quality control, alignment, duplicate marking, filtering, broad peak calling, and generation of counts per million (CPM)-normalized signal tracks. Reads were aligned to the Mus musculus reference genome (mm10), with the iGenomes GTF gene annotation file (UCSC) provided during pipeline execution for peak annotation. A read length of 100 bp was specified, which was used to estimate fragment size and inform background modeling during broad peak calling with MACS2. Regions in the mm10 ENCODE blacklist (v2) were excluded during execution using the --blacklist flag.

### CUT&RUN for histone modifications and LADs mapping

CUT&RUN was performed using the CUTANA ChIC/CUT&RUN Kit (EpiCypher) according to the manufacturer’s instructions. Two biological replicates were prepared for each antibody used. Briefly, P1 cardiomyocytes were isolated and purified as described above. Samples were divided into aliquots of 100,000 cells, with one aliquot from each sample reserved for the IgG control. Cells were washed and bound to activated Concanavalin A beads provided in the kit by incubating at room temperature for 10 minutes. Beads were magnetized, the supernatant was removed, and beads were resuspended in Antibody Buffer. Primary antibodies were added to each aliquot and incubated overnight at 4°C with gentle agitation. Antibodies used were: IgG control (0.5 μL; provided in kit), lamin-A/C (0.5 μL; Abcam, ab133256), H3K9me2 (0.5 μL; Diagenode, C15200154), H3K9me3 (0.5 μL; Active Motif, 39062), and H3K27ac (0.5 μL; EpiCypher, 13-0059). 2 μL of K-MetStat panel diluted 1:5 was added to control samples. The following day, beads were washed twice with 200 μL of Permeabilization Buffer and resuspended in 50 μL of Permeabilization Buffer containing 2.5 μL of pAG-MNase. After a 10-minute incubation at room temperature, beads were again washed twice with 200 μL of Permeabilization Buffer and resuspended in 50 μL of Permeabilization Buffer containing 1 μL of 100 mM calcium chloride. Reactions were incubated at 4°C for 2 hours with gentle agitation to allow digestion. Reactions were stopped by adding the kit-provided Stop Master Mix together with 1 μL of 1:5 diluted E. coli Spike-in DNA. Samples were incubated at 37°C for 10 minutes. The supernatant was collected, and DNA was purified using 119 μL of SPRIselect beads included in the kit, following the manufacturer’s instructions. Using ∼2 ng of DNA, sequencing libraries were prepared with the Takara ThruPLEX DNA-seq kit according to the manufacturer’s protocol, with 10 cycles of PCR amplification. Libraries were quantified with the Qubit dsDNA Quantification Assay Kit, and library quality was verified on the 2100 Bioanalyzer System with the High Sensitivity DNA Kit. Sequencing was performed on the Illumina NextSeq 1000 with 50 x 50 bp paired-end reads.

### Processing of CUT&RUN data

Samples were processed using the nf-core/cutandrun pipeline (v3.2.2) with Nextflow (v23.10.0). Reads were aligned to the mm10 Mus musculus reference genome, and mitochondrial reads were removed by including the --remove_mitochondrial_reads flag during pipeline execution. The pipeline performed adapter trimming, quality control, alignment, duplicate marking, and filtering. For Hidden Markov Modeling (HMM), replicate BAM files output by the pipeline were merged using samtools merge and converted to CPM-normalized bigWig files at 5 kb resolution. Histone and lamin CUT&RUN samples were normalized to their corresponding IgG control by running bigWigCompare with the --operation Log_2_ and --pseudocount 1 flags.

### Comparison of CUT&RUN Lamin-A LADs mapping among samples

For lamin CUT&RUN, additional z-score normalization was performed on the generated bigWig files to enable comparison between samples. To identify LAD regions, a two-state HMM was called using the depmixS4 R package (v1.4-0). Analyses were restricted to chromosomes 1-19, and the first 3 Mb of each chromosome was excluded to avoid boundary artifacts. Overlaps between lamin-B1 heterozygous and null LADs were identified, and LADs were classified as shared or unique. LADs present in the heterozygous track but absent from the null track were defined as “lost LADs.” For shared LADs, the fraction of each heterozygous LAD retained in the null sample was calculated by intersecting genomic ranges; LADs with greater than 10% loss were classified as LADs with edge loss.

### ChIP-seq processing of previously published datasets

Raw FASTQ files were downloaded from GSE155658^49^ for H3K9me1, H3K9me3, and H3K27me3. ChIP-seq data were processed using the nf-core/chipseq pipeline (v2.1.0) with Nextflow (v23.10.0). Replicate BAM files were merged using samtools merge and converted to CPM-normalized bigWig files at 5 kb resolution. Each ChIP-seq track was normalized to its corresponding input by running bigwigCompare with the --operation Log_2_ and --pseudocount 1 flags. For visualization of H3K4me3, bigWig files were generated using a 50 x50 bp bin size.

### Transcription factor binding motif analysis

H3K4me3 promoter peaks of differentially expressed genes were used as input to findMotifsGenome.pl from HOMER (v4.11), with the -size given flag to restrict the search to the provided peak regions. annotatePeaks.pl was then used to assign the top five enriched motifs to the H3K4me3 promoter peaks in both the upregulated and downregulated gene sets.

### Determination of HiLands chromatin states through HMM

Genome-wide signal tracks at 5 kb resolution for H3K9me2, H3K9me3, H3K27me3, H3K27ac, H3K4me1, H3K4me3, ATAC-seq, and lamin-associated domains (LADs) were compiled and processed as described above. ATAC-seq signals were Log_2_-transformed with a small pseudocount, while all other datasets were used as provided. Signal values across all tracks were capped at the range -3 to 3 to limit the influence of outliers. Analyses were restricted to autosomal chromosomes, and the first and last 3 Mb of each chromosome were excluded to reduce edge effects. Regions overlapping a combined blacklist (mm10 ENCODE blacklist v2 and low-signal regions) were also removed. All datasets were merged by genomic coordinates, and missing values were set to zero. PCA was performed on z-score-scaled signals from all tracks, and the first three principal components were retained for downstream analyses.

We then applied an HMM to infer chromatin states and define HiLands domains as previously described^55,61^, with emission probabilities modeled using a multivariate normal inverse Gaussian distribution. A six-state model was used to capture chromatin variation across the seven input tracks. Model parameters were estimated using the expectation-maximization (Baum-Welch) algorithm, with 20 random initializations to reduce convergence to local optima; the highest-likelihood solution was selected. Chromatin states were assigned using the Viterbi algorithm, and adjacent genomic windows sharing the same state were merged to generate continuous HiLands regions.

### Hi-C

#### 1. Cardiomyocyte crosslinking

P1 cardiomyocytes were isolated and purified as described above, and an aliquot of 10,000 cells per sample was used for Hi-C. Hi-C was performed using an optimized low-input protocol with carrier DNA; a comprehensive description of the methodology will be reported elsewhere and is available upon request. Cardiomyocytes were fixed with 2% PFA in PBS for 10 minutes at room temperature. PFA was quenched by adding glycine to a final concentration of 125 mM, followed by a 5-minute incubation at room temperature and a 15-minute incubation on ice. Cells were pelleted at 300 x g for 5 minutes at 4°C and washed with 1 mL of ice-cold PBS.

#### 2. Cell lysis and enzyme digestion

Cells were pelleted again under the same conditions and resuspended in 1.0 mL of Hi-C lysis buffer (10 mM Tris-HCl pH 8.0, 10 mM NaCl, 0.2% (volume/volume) IGEPAL CA-630 (NP-40), with 1X protease inhibitor cocktail). Cells were rotated for 30 minutes at 4°C and then pelleted at 2,000 x g for 8 minutes at 4°C. Pellets were washed with 500 μL of ice-cold 1.2X NEB3 buffer, pelleted under the same conditions, and resuspended in 40 μL of 1.2X NEB3 buffer. 1.2 μL of 10% SDS was added to each sample (final concentration 0.3%), and samples were incubated at 65°C for 10 minutes and then placed on ice. SDS was quenched by adding 4 μL of 20% Triton X-100 and gently mixing, followed by incubation at 37°C for 1 hour without agitation. 2.5 μL of DpnII restriction enzyme (25 units; NEB, R0543S, 10 units/μL) was then added to each sample and incubated for 90 minutes at 37°C with gentle mixing every 30 minutes. An additional 2.5 μL of DpnII was then added and incubated for another 90 minutes under the same conditions.

#### 3. Fill 5’overhang with biotin and proximity ligation

The 5’ overhangs generated by DpnII digestion were filled in with biotin by adding to each sample 0.5 μL of a 10 mM dCTP/dGTP/dTTP mix (Qiagen, 201912), 1.0 μL of 1 mM biotin-14 dATP (Jena Bioscience, NU-835-BIO14-S), and 1.5 μL of Klenow fragment (NEB, M0210L; 5 units/μL). Samples were incubated for 60 minutes at 37°C and then placed on ice. The blunt-ended DNA was then ligated by adding 50.0 μL of 10X T4 ligase buffer (NEB, B0202S), 5.0 μL of 100X BSA (NEB, B9000S), 390.8 μL of water, and 1.0 μL of T4 ligase (NEB, M0202S; 400 units/μL). Samples were incubated overnight at 16°C without agitation.

#### 4. Reversal of crosslinking and DNA purification

To reverse crosslinking, 2.5 μL of 20 mg/mL proteinase K (ThermoFisher, 25530049) was added to each sample, which was then incubated overnight at 65°C. The following day, an additional 2.5 μL of 20 mg/mL proteinase K was added, and samples were incubated for another 2 hours at 65°C. DNA was extracted by adding 500 μL of phenol/chloroform/isoamyl alcohol (25:24:1; Sigma, P2069), vortexing, and recovering the aqueous supernatant. Phenol/chloroform/isoamyl alcohol extraction was repeated once more, and the supernatant was transferred to a 2 mL Eppendorf tube. DNA was further purified using the Silica Bead DNA Extraction Kit (ThermoFisher, K0513) according to the manufacturer’s protocol. Briefly, 500 μL of supernatant was mixed with 1.5 mL of silica bead binding buffer and 10 μL of silica bead suspension. Samples were incubated at 55°C for 15 minutes with occasional mixing. Silica beads were then washed three times by pelleting and resuspending in 500 μL of the kit-provided ice-cold wash buffer. After the final wash, beads were completely air-dried. DNA was eluted by resuspending the beads in 45 μL of 10 mM Tris-HCl (pH 8.0) and incubating at 55°C for 5 minutes. Beads were pelleted, and the supernatant was transferred to a Spin-X tube filter (Sigma, CLS8162-24EA, or Costar, 8160) to remove residual beads. Tubes were centrifuged at 16,000 x g for 5 minutes at 4°C, and 40 μL of the filtered DNA solution was used for the de-biotinylation reaction.

#### 5. Removal of biotin from unligated ends

To remove biotin from un-ligated ends, 1.0 μL of 10 mg/mL BSA, 10.0 μL of 10X NEB2 buffer (NEB, B7002S), 12.5 μL of 10 mM dNTPs (Qiagen, 201912), 34.9 μL of distilled water, and 1.6 μL of T4 DNA polymerase (NEB; 3 units/μL) were added to each sample. The reaction was incubated at 20°C for 2 hours without agitation, and then stopped by adding 2 μL of 0.5 M EDTA. DNA was purified using the silica bead DNA extraction kit as described above, with the same buffer-to-sample ratios, but eluted in 50 μL of elution buffer.

#### 6. DNA sonication and end repair

50 μL of the DNA suspension was transferred to a microTUBE AFA Fiber Screw-Cap (Covaris, 520166) and sonicated using the Covaris M220 instrument with the M220 Holder-XTU holder (Covaris, 500488). Hi-C DNA was sheared for 90 seconds at 20°C using a peak incident power of 50 W, a duty cycle of 20%, and 200 cycles per burst. Sheared DNA was transferred to PCR tubes for end repair. End repair was performed by adding 10 μL of 10X phosphorylation reaction buffer (NEB, B0201S), 0.5 μL of T4 DNA polymerase (NEB, M0203S), 0.5 μL of T4 polynucleotide kinase (NEB, M0201S), 5 μL of 10 mM dNTPs (Qiagen, 201912), 15 μL of 10 mM dATP (NEB, N0440S), 2 μL of 10X-diluted Klenow fragment DNA polymerase I (NEB, M0210L), and 17 μL of distilled water. Samples were incubated at 37°C for 30 minutes without agitation.

#### 7. Streptavidin capture

100 μL of end-repaired Hi-C DNA was mixed with 100 μL of 2X BW buffer (10 mM Tris-HCl pH 7.5, 1 mM EDTA, 2 M NaCl) in preparation for streptavidin capture. 20 μL of Dynabeads M-280 Streptavidin (ThermoFisher, 11205D) was transferred to a low-adhesion 0.5 mL PCR tube (Sigma, CLS6530), and washed with 100 μL of 2X BW buffer. End-repaired DNA was added to the prewashed beads, mixed by gentle pipetting, and rotated at room temperature for 60 minutes. Samples were placed on a magnetic stand, and the supernatant was removed. Beads were resuspended in 1X BWT buffer (5 mM Tris-HCl pH 7.5, 0.5 mM EDTA, 1 M NaCl, 0.1% Triton X-100), magnetized, and the supernatant was removed; this wash was repeated once without changing tubes. Beads were then resuspended in 10 mM Tris-HCl (pH 8.0; Qiagen Buffer EB), magnetized, and the supernatant was removed; this wash was also repeated once. Finally, beads were resuspended in 16 μL of 10 mM Tris-HCl (pH 8.0).

#### 8. Library preparation and sequencing

A-tailing was performed on 16 μL of bead-bound Hi-C DNA by adding 2.5 μL of 10X NEB2 buffer (NEB, B7002S), 5.0 μL of 1 mM dATP (NEB, N0440S), and 1.5 μL of Klenow fragment (3’→5’ exo-) (NEB, M0212L; 5 units/μL), followed by incubation at 37°C for 30 minutes without agitation in a thermal cycler. Beads were then placed on a magnetic stand, the supernatant was removed, and beads were washed once with 100 μL of 1X BWT buffer. Beads were magnetized, the supernatant was removed, and beads were resuspended in 100 μL of elution buffer (10 mM Tris-HCl, pH 8.0); this wash was repeated for a total of two washes. After the final wash, the supernatant was removed, and beads were resuspended in 6.5 μL of elution buffer. Sequencing adaptors were ligated by adding 12.5 μL of Quick ligation reaction buffer (NEB, M2200L), 5.0 μL of 1:4-diluted NEBNext Adaptor (NEB, E7500S), and 1.0 μL of Quick T4 ligase (NEB, M2200L), followed by incubation at 20°C for 15 minutes. 1.5 μL of USER enzyme mix was then added, mixed by gentle pipetting, and incubated at 37°C for 15 minutes. Beads were magnetized and washed three times with 100 μL of 1X BWT buffer, changing tubes between washes, and twice with 100 μL of elution buffer without changing tubes. Beads were finally resuspended in 10 μL of elution buffer. PCR amplification reactions were assembled by adding 1.25 μL of Universal PCR primer (NEB, E7500S), 1.25 μL of the chosen Index primer (NEB, E7500S), and 12.5 μL of NEBNext Ultra II Q5 Master Mix (NEB, M0544S). PCR was performed with an initial denaturation at 98°C for 1 minute, followed by 12 cycles of 98°C for 10 seconds, 62°C for 75 seconds, and 72°C for 20 seconds, followed by a final extension at 65°C for 5 minutes and a hold at 10°C. Beads were magnetized, and the supernatant containing the PCR product was recovered. Finally, the 25 μL PCR product was purified by adding 20 μL of SPRIselect beads (Beckman Coulter; 0.8X volume ratio) and incubating at room temperature for 15 minutes. Beads were magnetized and washed twice with 200 μL of 80% ethanol while attached to the magnetic stand. Beads were air-dried and resuspended in 12 μL of elution buffer, magnetized, and the supernatant was collected. Final libraries were quantified with the Qubit dsDNA Quantification Assay Kit, and library quality was verified on the 2100 Bioanalyzer System with the High Sensitivity DNA Kit. Sequencing was performed on the Illumina NextSeq 1000 with 50 x 50 bp paired-end reads.

### Hi-C Analysis

#### 1. Hi-C data preprocessing

We used the 4DN Hi-C processing pipeline (https://github.com/4dn-dcic/docker-4dn-hic) for read alignment and filtering. Quality control of individual Hi-C libraries was performed independently using the HiCUP pipeline to assess the proportion of valid read pairs and overall library complexity.

#### 2. TAD boundary identification

To identify TAD boundaries, insulation scores were calculated using the fanc insulation command from the FAN-C package (v0.9.28) on the processed Hi-C data at 20 kb resolution. A 1 Mb window size was applied with the -w flag, as it produced insulation profiles that best corresponded to domain boundaries visible in the Hi-C contact maps. Insulation scores were smoothed using a rolling mean over three bins. Boundaries were defined as local minima in the smoothed insulation profile: for each genomic bin, the insulation score was compared to values within a ±8-bin window, and bins with values lower than all neighboring bins were called as boundaries. To retain only strong domain borders, boundaries were required to have insulation scores below -0.3.

#### 3. Hi-C contact visualization

For visualization and analysis of Hi-C contact maps, we applied a dynamic binning strategy to ICE-normalized contact matrices, similar to that used in our previous study^55^. To reduce noise in low-coverage regions, we implemented an adaptive smoothing approach: for each bin pair, if the observed raw contact count met a minimum threshold of 20 reads, the corresponding scaled ICE value was retained; otherwise, the bin pair was iteratively expanded symmetrically in both genomic dimensions until the cumulative raw contact count within the expanded region reached the threshold, or until the matrix boundaries were reached. The contact value for the original bin pair was then defined as the mean of the scaled ICE values within this expanded window. This dynamic binning procedure was applied independently to each condition, producing smoothed contact matrices for both control and mutant samples. For comparative analysis, Log_2_ fold-change matrices were computed by taking the Log_2_ ratio of the dynamically binned, scaled ICE matrices between conditions, using a small symmetric pseudocount to avoid division by zero. This approach reduces sampling noise in sparse regions while preserving high-resolution features in well-covered regions, enabling robust comparison between conditions.

To further visualize locus-specific interaction changes, pseudo-4C profiles were generated from the Log_2_FC matrices. For a given gene of interest, the genomic bin corresponding to its promoter was selected as an anchor point. Interaction frequencies between this anchor bin and all other bins across the region of interest were extracted directly from the Log_2_FC matrix, producing a one-dimensional interaction profile. To reduce local noise and enhance interpretability, the signal was smoothed using a rolling mean with a fixed window size, followed by linear interpolation to handle missing values.

#### 4. A/B compartment analysis

Compartment scores were calculated using the previously described C-score tool^58^. Raw deduplicated contact pairs generated by the 4DN processing pipeline for each genotype were used as input, and scores were calculated at 5 kb resolution. Interactions spanning less than 1 Mb were excluded to remove contacts more strongly associated with TAD-level interaction dynamics. C-score tracks were then compared with our LAD maps to identify A and B compartment regions. To ensure consistency with LAD assignments, chromosome-wide C-scores were multiplied by -1 where necessary to invert the C-scores, such that B compartments corresponded to positive values and A compartments corresponded to negative values across the genome.

#### 5. Hi-C interaction changes between HiLands and gene promoters

Hi-C interaction data were processed from tab-delimited files containing paired genomic bin coordinates and contact counts, retaining only intrachromosomal interactions on chromosomes 1-19. Chromatin interaction analysis focused on promoter-centered contacts defined using H3K4me3-marked promoter annotations, with promoter regions extended ±5 kb and additional upstream and downstream flanking regions defined as 5 Mb windows relative to gene boundaries. Chromatin states were assigned to Hi-C bins by overlapping interaction anchors with HiLands genomic state intervals, enabling annotation of each contact with state information for both interacting bins. Genes were stratified into upregulated, downregulated, and non-differentially expressed groups based on adjusted p-values and log_2_FC thresholds from the differential expression analysis, and promoter and flanking regions were subset accordingly. Hi-C contact counts were normalized across datasets by scaling each sample to a common total contact depth derived from the mean library size across all replicates. Promoter-flank interactions were extracted by retaining contacts in which one anchor overlapped a promoter region and the other overlapped either an upstream or downstream flanking region, considering both interaction orientations. For each gene, scaled contact counts were aggregated across interactions and summarized by flanking chromatin state using HiLands annotations, producing gene-level contact profiles stratified by state. Differential interaction effects were quantified as Log_2_FCs between lamin-B1 knockout and control mean contact frequencies per gene and chromatin state. Statistical significance of differences between gene classes (upregulated vs non-DE, downregulated vs. non-DE) was assessed within each chromatin state using Wilcoxon rank-sum tests.

## QUANTIFICATION AND STATISTICAL ANALYSIS

Quantification methods, including the number of biological replicates, sample sizes, and statistical tests used, are described in the corresponding figure legends and methods sections. Data distributions were not formally tested for normality. No statistical methods were used to predetermine sample size, and there was no blinded analysis.

## Supplemental Figure

**Figure S1.**
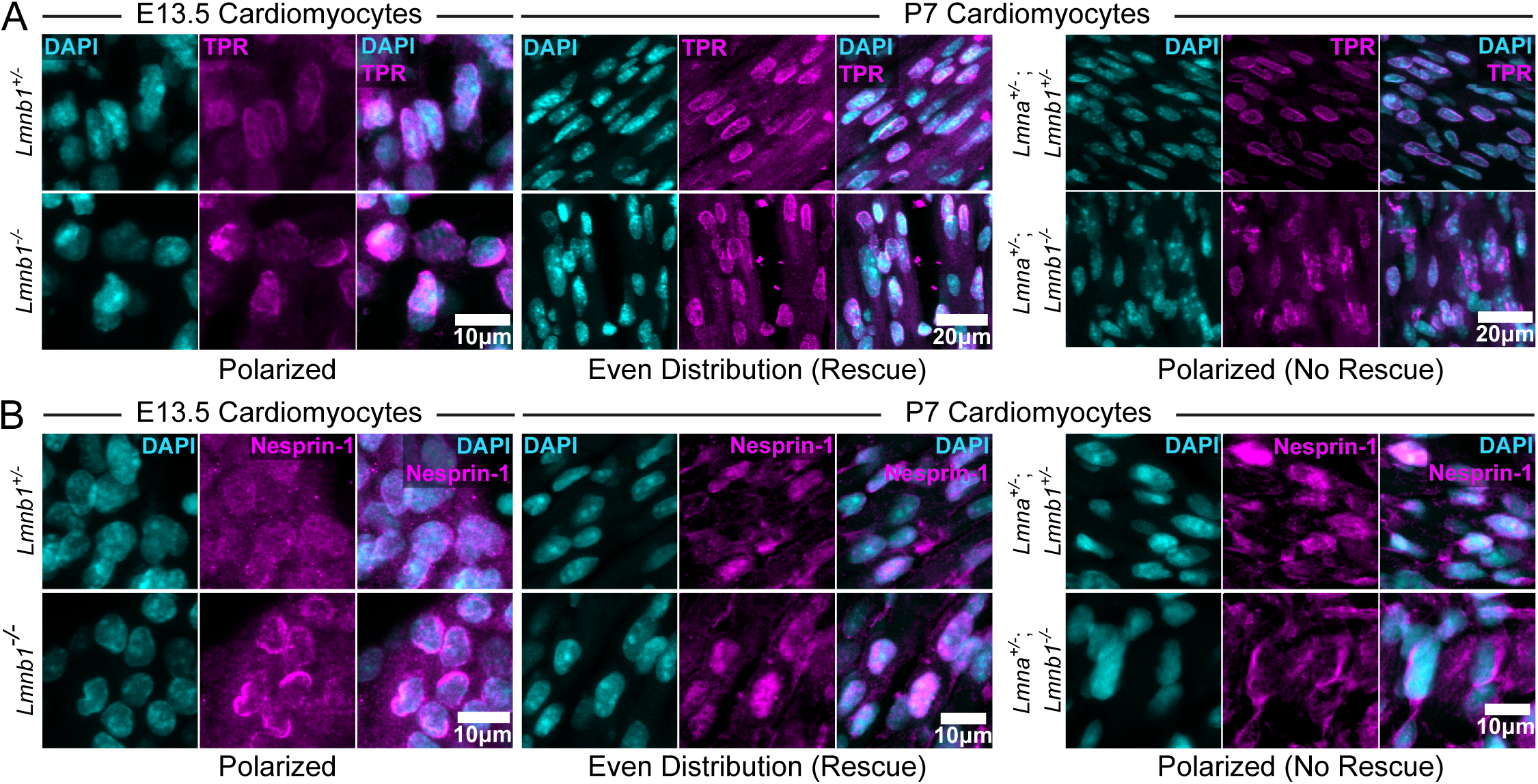
Postnatal upregulation of lamin-A rescues distribution of NPC and LINC complex in *Lmnb1*^-/-^ but not *Lmna*^+/-^; *Lmnb1*^-/-^ cardiomyocytes. (A, B) Representative confocal images of heart tissue sections immunostained for the nuclear pore complex component TPR (A, magenta) or the LINC complex component Nesprin-1 (B, magenta) and DAPI (cyan) at E13.5 (left panels) and P7 (center and right panels). The scale bars are as indicated. N= 3 biological replicates per stain and genotype.

**Figure S2.**
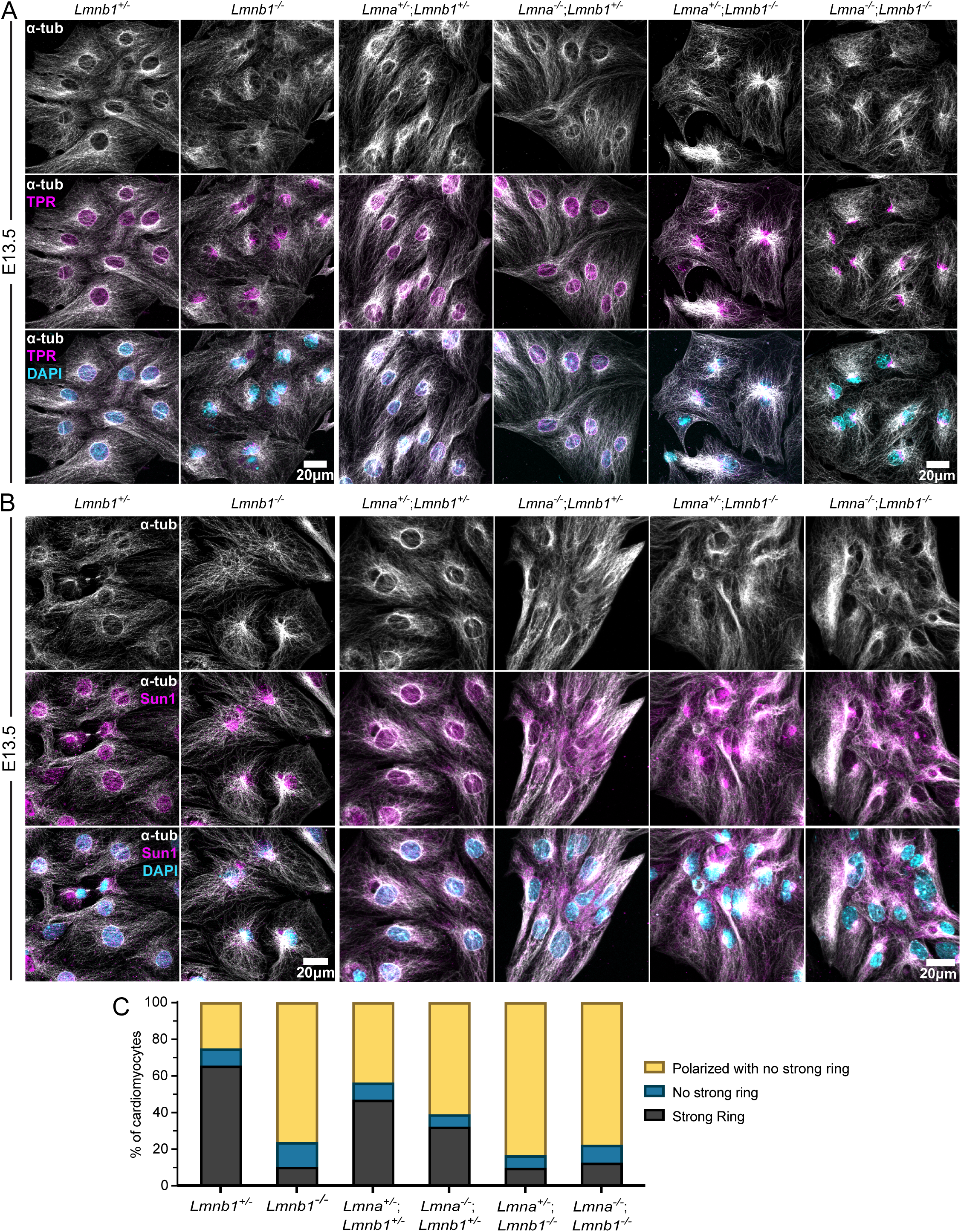
Loss of lamin-B1 promotes astral microtubule arrays and disrupts the perinuclear microtubule cage in E13.5 cardiomyocytes. (A,B) Representative confocal images of cultured E13.5 cardiomyocytes immunostained for α-tubulin (α-tub, white), DAPI (cyan), and either the NPC marker TPR (magenta) (A) or the LINC complex component Sun1 (magenta) (B) across the indicated genotypes. Scale bar, 10 μm. (C) Stacked bar plot showing quantification of microtubule organization categories in E13.5 cardiomyocytes across the indicated genotypes. Bars represent the percentage of cardiomyocytes displaying each microtubule phenotype: strong perinuclear ring (black), astral array with no strong ring (blue), or polarized astral array (yellow). Data represents mean and n= 3 biological replicates.

**Figure S3.**
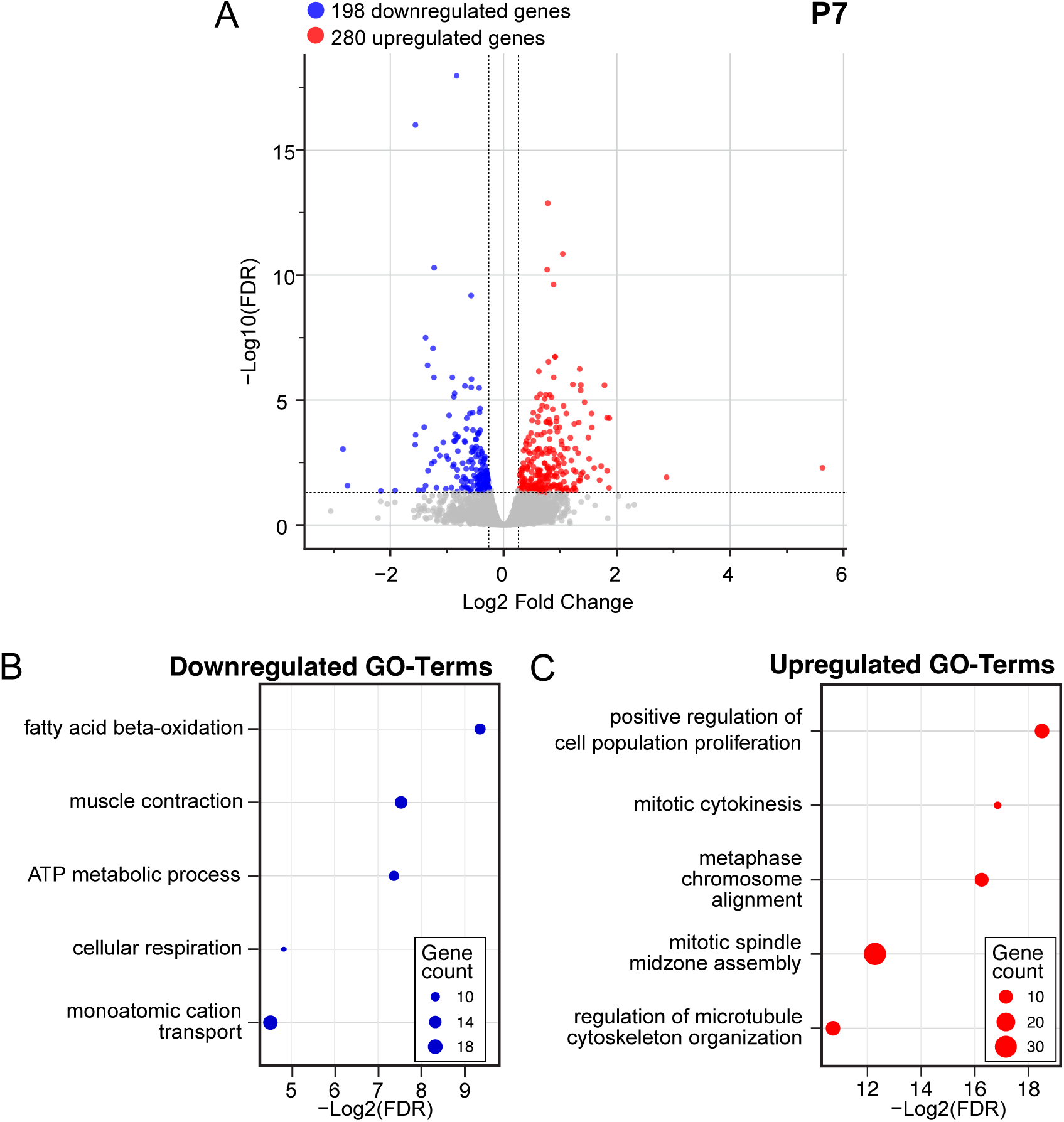
Bulk RNA-seq of P7 *Lmnb1^-/-^* hearts reveals downregulation of cardiomyocyte maturation genes and upregulation of cell cycle genes. (A) Volcano plot of differentially expressed genes in *Lmnb1^-/-^* versus *Lmnb1^+/-^* P7 whole hearts by bulk RNA-seq. Downregulated genes and upregulated genes are shown in blue and red, respectively. The dashed lines indicate significance thresholds (|Log_2_FC| > 0.26; p < 0.05). (B, C) Selected GO-terms enriched among downregulated (B) and upregulated (C) genes.

**Figure S4.**
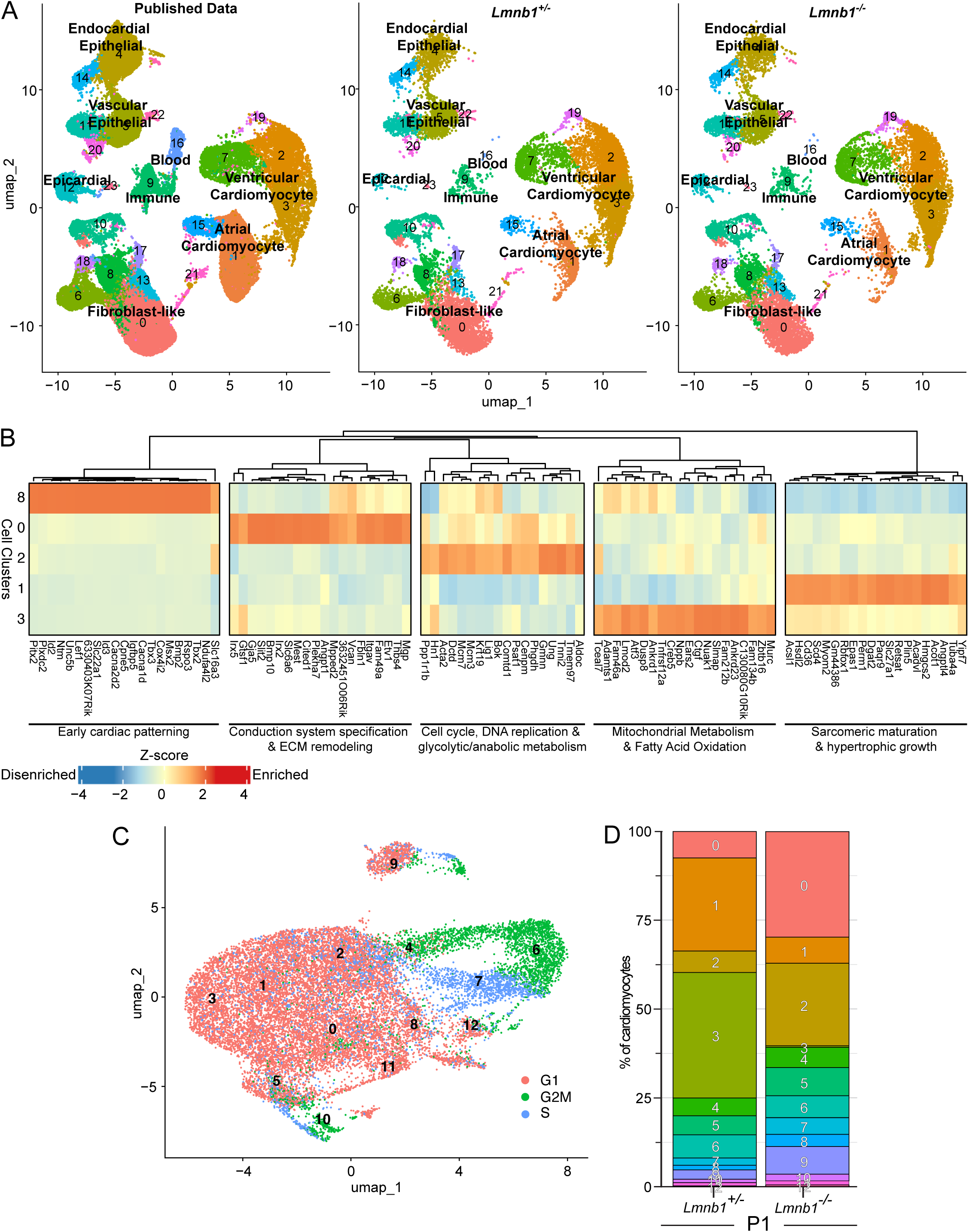
Identification of major cell populations in the heart and characterization of ventricular cardiomyocyte clusters. (A) UMAPs of the integrated scRNA-seq dataset showing major cardiac cell populations, displayed separately for previously published data (left), *Lmnb1^+/-^* (middle), and *Lmnb1^-/-^*(right) datasets. Major cell populations are denoted, including ventricular cardiomyocytes, atrial cardiomyocytes, fibroblast-like cells, epicardial cells, endocardial/epithelial cells, vascular epithelial cells, blood cells, and immune cells. (B) Heatmap showing scaled expression (z-score) of top marker genes across five compact ventricular cardiomyocyte clusters (clusters 0, 1, 2, 3, and 8) in previously published datasets, used to assign developmental stage identities. (C) UMAP of ventricular cardiomyocytes colored by cell cycle phase (G1, red; G2M, green; S, blue) inferred with Seurat’s CellCycleScoring package. (D) Stacked bar plots showing the percentage of ventricular cardiomyocytes from each cluster in the *Lmnb1^+/-^*and *Lmnb1^-/-^* P1 datasets.

**Figure S5.**
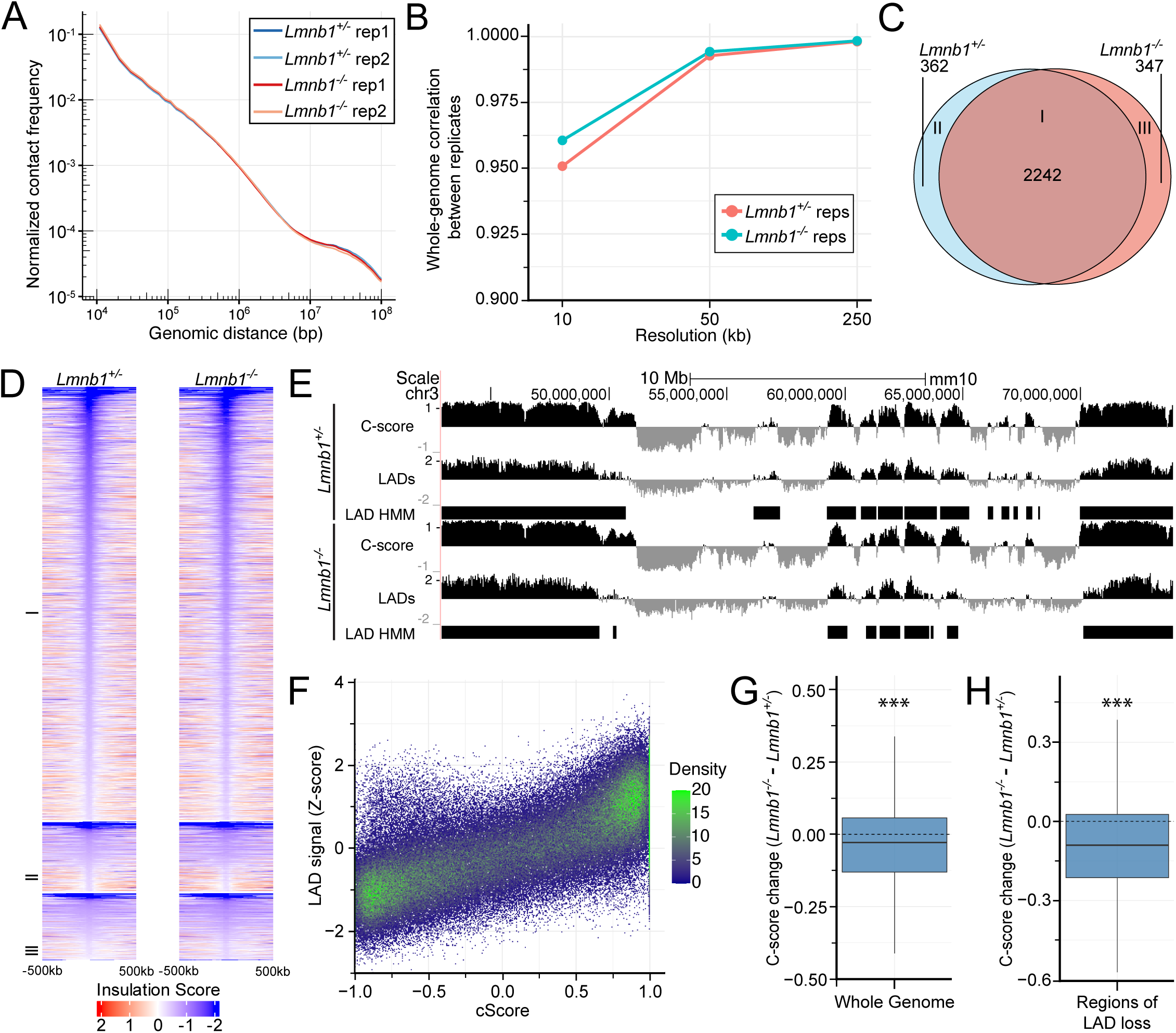
Hi-C quality control and compartment analyses in P1 cardiomyocytes. (A) Normalized contact frequency as a function of genomic distance for all Hi-C replicates, showing high consistency between *Lmnb1^+/-^* replicates (dark/light blue) and *Lmnb1^-/-^* replicates (red/orange). (B) Whole-genome correlation between biological replicates across increasing Hi-C resolutions (10–250 kb) for *Lmnb1^+/-^* (pink) and *Lmnb1^-/-^* (teal) samples. (C) Venn diagram showing the number of TAD boundaries shared between and unique to *Lmnb1^+/-^* and *Lmnb1^-/-^* cardiomyocytes. (D) Heatmaps showing insulation scores in ±500 kb around TAD boundaries at 20 kb resolution in *Lmnb1^+/-^* and *Lmnb1^-/-^* P1 cardiomyocytes. Groups I, II, and III correspond to TAD boundaries shared between genotypes, unique to *Lmnb1^+/-^*, and unique to *Lmnb1^-/-^*, respectively, as defined in (C). (E) Genome browser tracks showing C-scores and LADs mapping of lamin-A CUT&RUN z-score signal with HMM-defined LAD calls for a representative region of chromosome 3 in *Lmnb1^+/-^*(top) and *Lmnb1^-/-^* (bottom) cardiomyocytes. Positive C-scores indicate B-compartment and negative C-scores indicate A compartment regions. (F) Density scatter plot showing the correlation between lamin-A CUT&RUN z-score signal and C-score across the genome, with color indicating point density. (G) Box plot showing the genome-wide change in C-score (*Lmnb1^-/-^* - *Lmnb1^+/-^*) across all genomic bins. (***p < 0.001, Wilcoxon rank-sum test). (H) Box plot showing the change in C-score (*Lmnb1^-/-^* - *Lmnb1^+/-^*) restricted to regions of LAD loss. (***p < 0.001, Wilcoxon rank-sum test).

**Figure S6.**
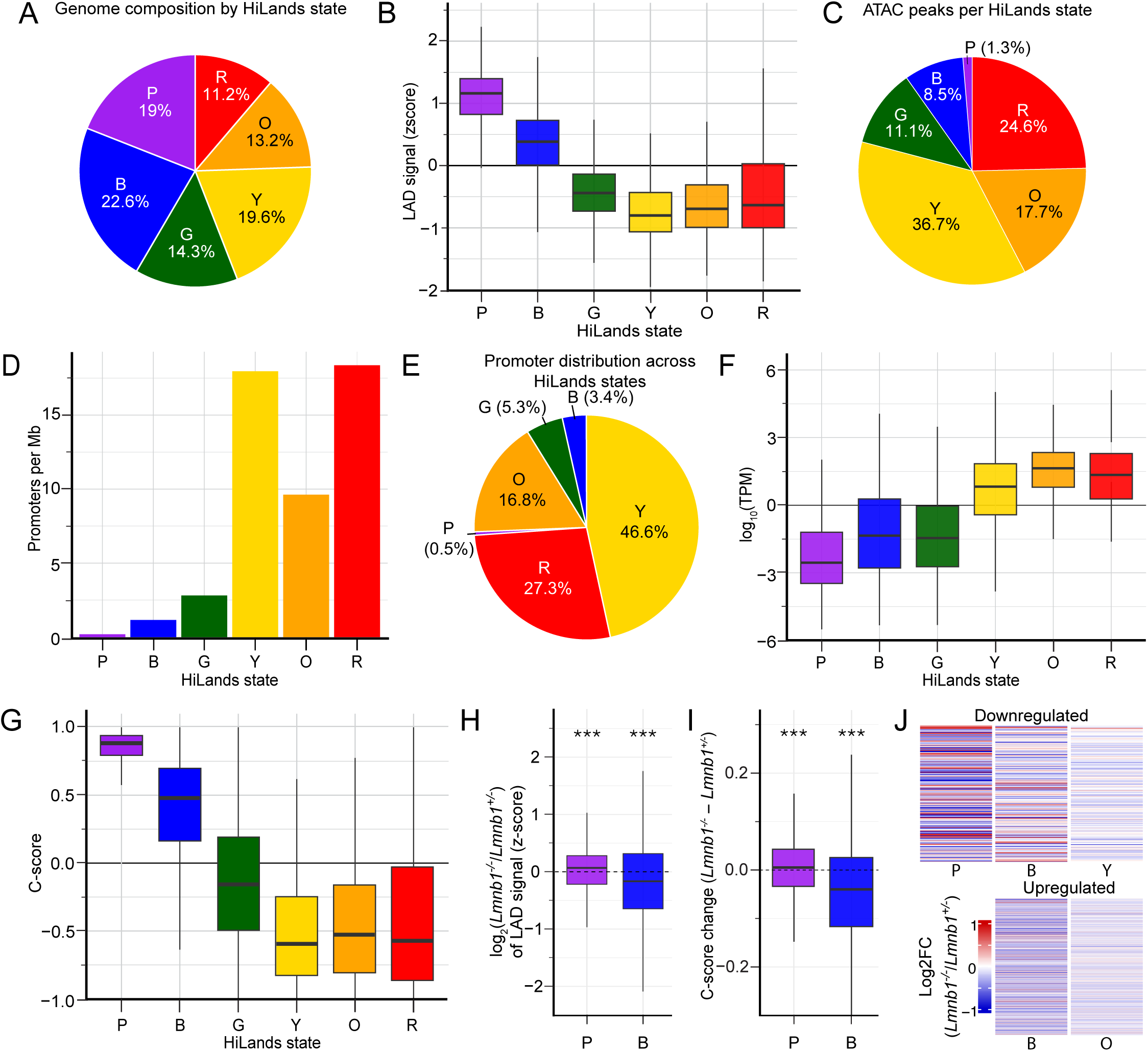
HiLands characterization in *Lmnb1^+/-^* P1 cardiomyocytes. (A) Pie chart showing the genome composition by HiLands state in wild-type P1 cardiomyocytes. Percentages indicate the fraction of the genome occupied by each state. (B) Box plots showing LADs mapping with lamin-A CUT&RUN z-score signal of bins across each HiLands state. (C) Pie chart showing the distribution of ATAC-seq peaks across HiLands states, with HiLands-Y, -O, and -R containing the majority of accessible chromatin regions. (D) Bar plot showing promoter density (promoters per Mb) across HiLands states. (E) Pie chart showing the overall distribution of gene promoters across HiLands states. (F) Box plots showing gene expression levels (log10 TPM) across HiLands states in P1 *Lmnb1^+/-^* compact cardiomyocytes from scRNA-seq data. (G) Box plots showing the C-score for each HiLands state bin in P1 cardiomyocytes. (H) Box plots showing the Log_2_FC of lamin-A CUT&RUN z-score signal within HiLands-P and HiLands-B regions in *Lmnb1^-/-^*relative to *Lmnb1^+/-^* cardiomyocytes. Box plots depict median and interquartile range. (***p < 0.001, Wilcoxon rank-sum test). (I) Box plots showing the genome-wide C-score change (*Lmnb1^-/-^* - *Lmnb1^+/-^*) within HiLands-P and HiLands-B regions. Box plots depict median and interquartile range. (***p < 0.001, Wilcoxon rank-sum test). (J) Heatmaps showing the mean Log_2_FC in 3D chromatin interactions between promoters of upregulated and downregulated genes and each HiLands state within ±5 Mb of individual promoters in *Lmnb1^-/-^* relative to *Lmnb1^+/-^* cardiomyocytes. Downregulated gene promoters show significantly increased interactions with HiLands-P (p = 0.0002) and -B (p = 0.003), as well as decreased interactions with HiLands-Y (p = 0.003) compared to non-dysregulated genes. Upregulated gene promoters show significantly reduced interactions with HiLands-B (p = 0.01), Hilands-O (p = 0.047) compared to non-dysregulated genes. (Wilcoxon rank-sum test).

## Supplemental Tables

**Table 1:** Dysregulated lists from P7 Bulk RNA-seq with downregulated genes (Sheet 1) and upregulated genes (Sheet 3) for *Lmnb1^-/-^*cardiomyocytes, as well as associated GO-terms for downregulated (Sheet 2) and upregulated (Sheet 4) lists. Related to Figure S3.

**Table 2:** Dysregulated lists from P1 scRNA-seq with downregulated genes (Sheet 1) and upregulated genes (Sheet 3) for *Lmnb1^-/-^*cardiomyocytes, as well as associated GO-terms for downregulated (Sheet 2) and upregulated (Sheet 4) lists. Related to Figure 4.

**Table 3:** Statistics of our lamin-B1 heterozygote and lamin-B1 knockout Hi-C Libraries. Related to Figure 5 and Figure S5.

**Table 4:** List of downregulated genes in *Lmnb1^-/-^*cardiomyocytes from P1 scRNA-seq that are explained by either changes in promoter interactions with either HiLands-B or -P (Sheet 1) or genes with a Jun binding motif (Sheet 2). List of upregulated genes in *Lmnb1^-/-^* cardiomyocytes from P1 scRNA-seq that are explained by either changes in promoter interactions with either a HiLands-B (Sheet 3) or genes with a E2f3 binding motif (Sheet 4). Related to Figure 6, 7, and S6.

**Table 5:** Homer motif output of downregulated gene promoters in *Lmnb1^-/-^* cardiomyocytes from P1 scRNA-seq (Sheet1) and associated list of downregulated genes with binding motifs for Sp1, Mef2d, E2f, Nf1, and Banp (Sheet 2). Homer output of upregulated gene promoters in *Lmnb1^-/-^*cardiomyocytes from P1 scRNA-seq (Sheet 3) and associated list of upregulated genes with binding motifs for Sp1, Mef2d, E2f, Nf1, and Banp (Sheet 4). Related to Figure 7.

**Table 6:** Primer list used for genotyping. Related to STAR Methods.

## Notes

### Competing Interest Statement

The authors have declared no competing interest.

### Summary of Updates

Additional edits were made to figures and text for grammar and additional clarity of findings.

